# TSignal: A transformer model for signal peptide prediction

**DOI:** 10.1101/2022.06.02.493958

**Authors:** Alexandru Dumitrescu, Emmi Jokinen, Juho Kellosalo, Ville Paavilainen, Harri Lähdesmäki

## Abstract

Signal peptides are short amino acid segments present at the N-terminus of newly synthesized proteins that facilitate protein translocation into the lumen of the endoplasmic reticulum, after which they are cleaved off. Specific regions of signal peptides influence the efficiency of protein translocation, and small changes in their primary structure can abolish protein secretion altogether. The lack of conserved motifs across signal peptides, sensitivity to mutations, and variability in the length of the peptides, make signal peptide prediction a challenging task that has been extensively pursued over the years. We introduce TSignal, a deep transformer-based neural network architecture that utilizes BERT language models (LMs) and dot-product attention techniques. TSignal predicts the presence of signal peptides (SPs) and the cleavage site between the SP and the translocated mature protein. We show improved accuracy in terms of cleavage site and SP presence prediction for most of the SP types and organism groups. We further illustrate that our fully data-driven trained model identifies useful biological information on heterogeneous test sequences.

## 1 Introduction

Signal peptides (SPs) are short amino acid chains found at the N-terminus of newly synthesized proteins. Their role is to facilitate the translocation of proteins, after which they are cleaved off from the mature protein by signal peptidases (SPases). SPs may direct proteins to the secretory (Sec) pathway (in all organisms) or twin-arginine translocation pathway (TAT), which is found only in prokaryotes and in plant chloroplasts. Proteins enter the Sec pathway in an unfolded state, while those going through the Tat pathway fold before the translocation.

Almost all SPs overall contain a tripartite structure with, generally positively charged, N-region, H-region (hydrophobic region) and a cleavage site-containing C-terminal region. In SPs cleaved by SPase I the cleavage site is preceded by a generally polar C-region, while SPs cleaved by SPase II have a three residue lipobox instead of the C region (Owji *et al.*, 2018) (see Fig. 1) and also a cysteine residue after the cleavage site (Tokunaga *et al.*, 1982). SPs processed by SPase IV do not have a tripartite structure, but instead contain a translocation-mediating basic region. Each of the aforementioned SP regions, which can vary in length and residue composition, dynamically interact with various components of the Sec or Tat machinery in order to facilitate protein translocation (Owji *et al.*, 2018). While SPs have recognizable regions, they lack clear consensus motifs. Consequently, the exact sequence properties of functional SPs have not been determined. This makes SP prediction challenging, which is evident in the problems of identifying translocation-abolishing point mutations (Liu *et al.*, 2012; Rajpar *et al.,* 2002).

**Figure 1:**
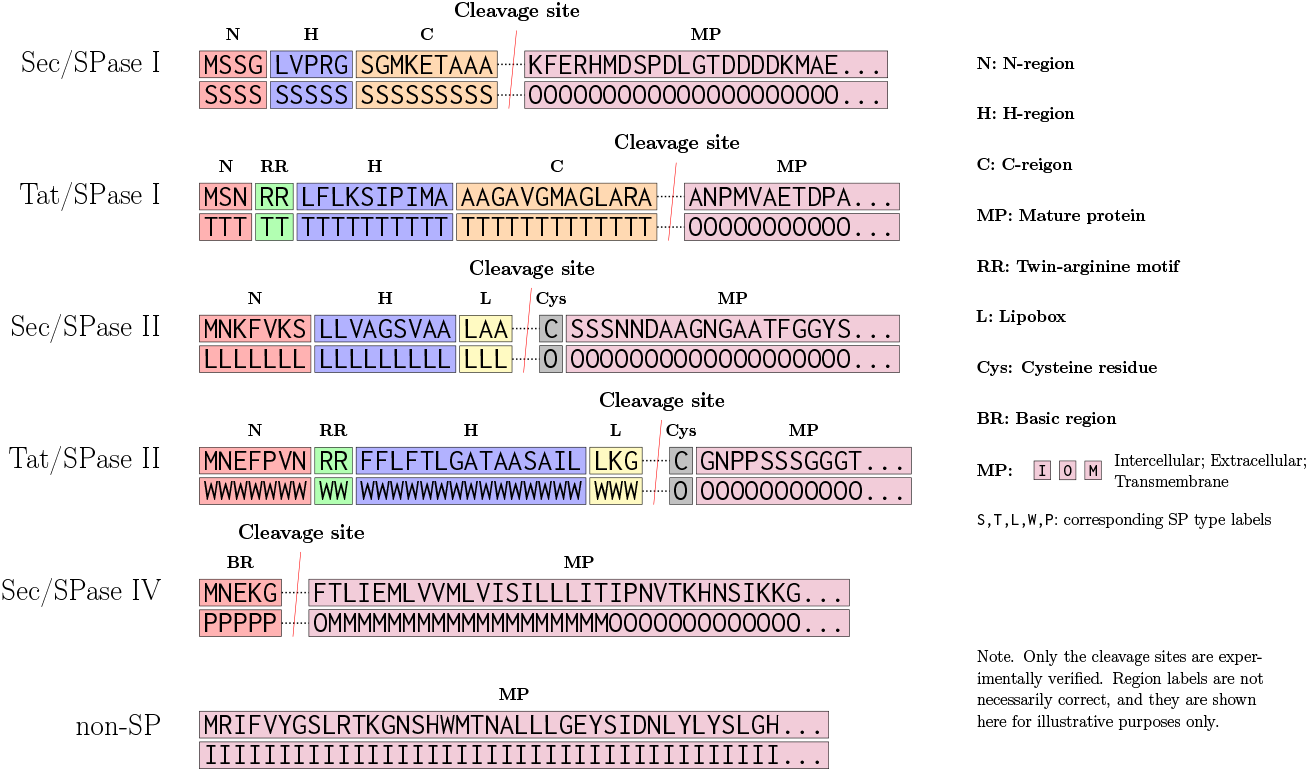
Examples of input sequences and associated labels for TSignal. Secretory path directed SPs cleaved by SPaseI have all three of the N, H and C subregions. Tat directed proteins present the twin-arginine (RR) motif in their SPs and the SP also has its N and H regions delimited by the RR motif. SPase II cleaved proteins do not present a proper C region but instead have three amino acids belonging to a region called lipobox (L), which is always followed by a Cys residue. SPase IV cleaved SPs only have a small basic region (BR) at the N-terminus. Amino acid sequence of an intracellular protein that does not contain a signal peptide is included as a reference.

Many of the previous machine learning approaches for signal peptide detection and cleavage site prediction rely on different types of hidden Markov models (HMMs) (Reynolds *et al.*, 2008; Viklund *et al.*, 2008; Tsirigos *et al.*, 2015; Käll *et al.*, 2004). Deep learning approaches have also been employed for the feature representations of the sequence residues. However, the final prediction is still carried out by structured prediction algorithms, using the deep residue representations as the inputs for conditional random fields (CRFs) (Zhang *et al.*, 2020; Savojardo *et al.*, 2018; Teufel *et al.*, 2022). Non machine-learning methods have also been employed. Homology-based search algorithms are used to detect the presence of SPs and report putative cleavage sites by aligning the queries to annotated sequences (Frank and Sippl, 2008; Wishart *et al.*, 2007). Despite several approaches to design and optimize SP prediction methods, no single approach provides robust SP identification. This difficulty is highlighted by the fact that protein database Uniprot relies on four separate SP prediction programs for SP assignment (Consortium, 2019).

An important aspect is that both HMMs and CRFs rely on a matrix of transition probabilities between consecutive amino acids. Using this matrix, prior information about the structure of SPs can be hard-coded into a model by constraining the transitions known to be biologically impossible to have zero probability. As a concrete example, state transition matrices can be restricted to produce only contiguous SP predictions and ensure they always start at the N-terminus of the sequence (Owji *et al.*, 2018), while cleavage sites for SPase II cleaved proteins can additionally be constrained to always be followed by a cysteine residue (Tokunaga *et al.*, 1982). While incorporating such inductive bias is generally found useful for classical HMM and CRF models, we do not enforce any such prior information. Here, we developed a new data-driven prediction method, TSignal, that uses transformer-based architectures both for the residue representation, as well as the prediction network. We utilize rich representations of amino acid sequences obtained from a BERT LM, and further train the BERT model together with self-attentive prediction methods. Using data from large databases of known signal peptides we demonstrate that our model achieves state-of-the-art performance compared to previous best approaches, including SignalP 6.0 (Teufel *et al.*, 2022).

## 2 Materials and methods

A protein is defined by its amino acid sequence **a** = (*a*_1_*a*_2_…*a_n_*) together with an associated label for each amino acid residue **y** = (*y*_1_*y*_2_…*y_n_*)· We consider the following eight labels, 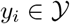 = {Sec/SPase I, Sec/SPase II, Sec/SPase IV, Tat/SPase I, Tat/SPase II, intracellular, transmembrane, extracellular}. The model distinguishes between five different SP types: secretory pathway directed peptides cleaved by SPase I, II and IV, and Tat pathway directed peptides cleaved by SPase I and II. Our model solves the cleavage site prediction task by predicting sequences of labels. The model predicts residue *α_i_* to be part of a signal peptide if its predicted label *ŷ_i_* corresponds to one of the five SP types we train for. The predicted SP type is inferred from the label *ŷ*_1_ associated with residue *a*_1_, while the cleavage site is determined by the first residue *α_c_* that is predicted to have one of the three non-SP labels *ŷ_c_* ∈ {intracellular, transmembrane, extracellular} following a sequence of predicted SP labels. This denotes that the cleavage site is located between residues *α*_*c*-1_ and *a_c_*. Additionally, each sequence **a** originates from one of the four organism groups *g* ∈ {eukarya, gn-bacteria, gp-bacteria, archaea}, corresponding to eukaryotes, gram-negative bacteria, gram-positive bacteria and archaea. Thus, each data item can be represented by a triplet (**a, y, *g***).

The structures of the SP types predicted by TSignal are shown in Figure 1. Although we do not explicitly utilize this information, we show that the model can intrinsically learn this type of structural information and generalize on diverse protein sequences.

### 2.1 Transformer models

Contrary to previous approaches that use HMMs or CRFs, our transformer model does not have any hard-coded knowledge of signal peptide structures. Instead, TSignal builds on the transformer model architecture which was initially developed for sequence translation tasks in the natural language field (Vaswani *et al.*, 2017). We use a contextual protein embedding model trained on 216 million protein sequences called ProtBERT (Elnaggar *et al.*, 2021). ProtBERT is a character-level adaptation of the auto-encoder BERT LM that is trained only on the masked-token prediction task. We use this model to retrieve the 1024 dimensional representations of all residues in each amino acid sequence, resulting in an *R*^1024×*N*^ representation for each sequence, with *N* being the maximum sequence length. To further adjust the ProtBERT model for signal peptide sequences, we integrate the BERT model as part of our model and train it together with the sequence prediction model, using a similar approach to SignalP 6.0. The 1024 × *N* sequence representations retrieved by ProtBERT are used as keys and values for the transformer decoder, along with the label queries in the multi-head attention blocks of the decoder. This way, we effectively train a sequence-to-sequence transformer architecture, as described by Vaswani *et al.* (2017). TSignal model architecture is shown in Figure 2 and described in more detail below.

**Figure 2:**
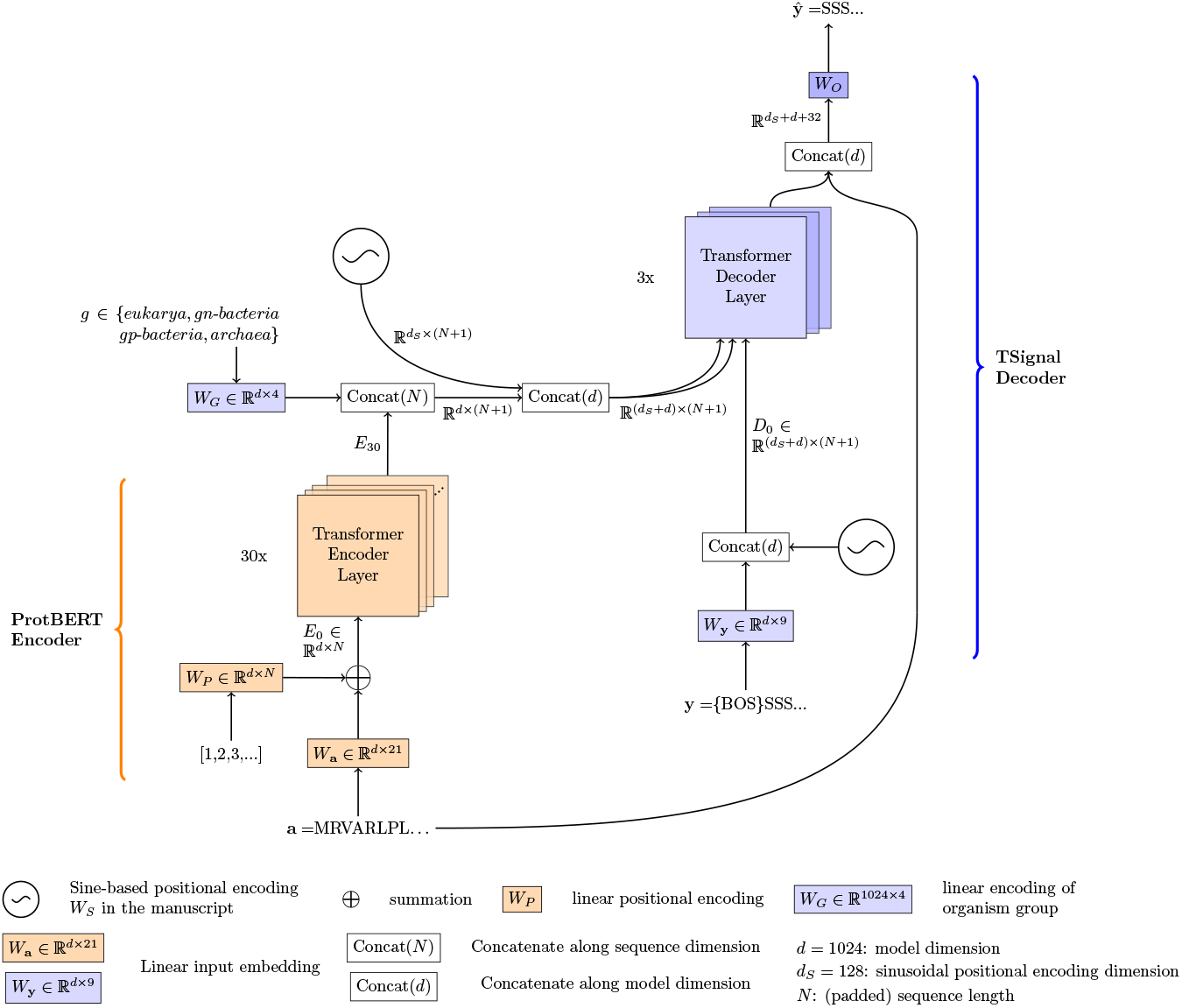
TSignal architecture consists of ProtBERT encoder and multi-head attention based transformer decoder. Initial embedding involves a dense representation and a linear positional encoding. ProtBERT embeddings are concatenated with organism group and positional representations prior to using them as key and value vectors in the decoder, together with query vector from dense representations of the sequence labels. See Section 2.1 for an in-depth description of the TSignal model.

#### 2.1.1 Encoder model

Transformer encoders use initial embedding layers to map tokens from one-hot vectors to dense representations. The positional encoding and multi-head attention mechanism are then able to extract contextual vector representations of these input tokens.

##### Initial embedding

Each amino acid *a_i_* of a sequence **a** = (*a*_1_*a*_2_ … *a_n_*) is initially one-hot encoded into 21-dimensional standard unit column vector **c**^(*i*)^ (from 20 unique amino acids and one additional padding token). For the whole sequence **a** this results in a binary one-hot encoded matrix *A* = (**c**^(1)^,…, **c**^(*n*)^) of size 21-by-*n*. If *n* is smaller than the maximum sequence length *N* in a mini-batch, then A is padded with *N* – *n* one-hot vectors corresponding to an additional padding token, forming a matrix of size 21-by-*N*. The initial embedding involves a linear transformation that maps each one-hot encoded amino acid to a dense d-dimensional^1^ representation using a matrix *W*_a_ ∈ *R*^*d*×21^. Collectively for the whole sequence **a** this can be written as a matrix multiplication *I* = *W*_a_*A* ∈ *R*^*d×N*^.

##### Positional encoding

The encoder model uses a linear positional encoding^2^. Since the amino acid indices will always be ordered from 1 to *N* (we assume the left-most amino acid is always the first N-terminus residue), we can directly define our positional representation for the amino acids as *W_P_* ∈ *R^d×N^*. With the above definitions, the initial positionally encoded embedding matrix for a sequence is:

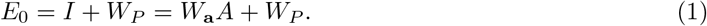

##### Transformer encoder

Representation 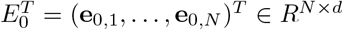 from Eq. (1) is then passed to multiple transformer block layers, where **e**_0,*i*_ is the d-dimensional initial representation of amino acid residue at position *i*. The core idea of transformer blocks is to process sequential information using only attention mechanisms, without any recurrent neural networks. In particular, transformers use dot-product attention. In one attention head *h* of layer *l*, attention weights 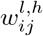 for a query residue *i* and key residue *j* are computed using the *d*-dimensional residue representations from the previous layers **e**_*l*-1,*i*_ and **e**_*l*-1,*j*_. This is done by first mapping the two vectors **e**_*l*-*1,i*_ and **e**_*l*-1,*j*_ into a dot-product suitable space and then computing their dot product

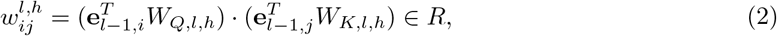

where · denote the vector dot-product and *W_Q,l,h_, W_K,l,h_* ∈ *R^d×d/H^* are the query and key linear layers of attention head *h* at layer *l,* and *H* is the total number of heads per layer (model dimension *d* is usually chosen to be exactly divisible by *H*). For a given query residue i, the attention weights 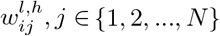 are normalized with softmax transformation, denoted as 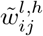. These attention weights are then used to compute a weighted average of the vectors formed by a third linear mapping (called value matrix) *W_V,h,l_* ∈ *R^d×d/H^* of the intermediate vectors that map the previous layer representations **e**_*l*-1,*j*_ into value vectors

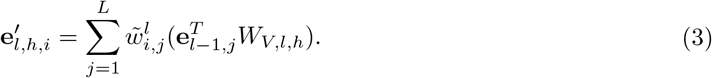

Multiple such attention “heads” are used in each layer, allowing the model to attend, at each step, to various parts of the sequences using different pairs of linear mappings (*W_Q,l,h_, W_K,l,h_*), where *h* ∈ {1,…, *H*} and *l* ∈ {1,…, *L*}. Lets define the key, query and value matrices computed by an attention head at layer *l* as

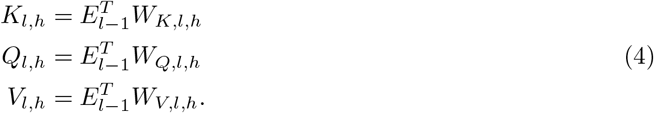

Note that the product 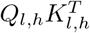 gives all the unnormalized weights 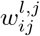 from Equation (2). The weighted average from Equation (3) for head *h* can be written compactly, for the whole sequence as

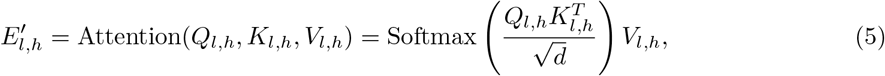

and the resulting sequence representations 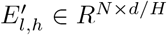 from all heads *h* ∈ {1,…, *H*} are stacked to obtain the intermediate matrix representation 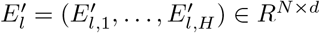.

Finally, the intermediate representation 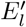 given by the multi-head attention at layer *l* is passed to a two-layered feed-forward network of the form 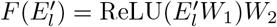. Usually, *W*_1_ is chosen to expand the model’s dimension from *d* to *e_d_, W*_1_ ∈ *R^d×e_d_^*, with *e_d_* being some expanding dimension, *e_d_* > *d*, and *W*_2_ ∈ *R^e_d_×d^* map the vectors back to dimension d (the network applies this transformation to each residue, individually). Skip connection and layer normalization layers are also added from *E*_*l*-1_ to 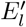 and from 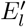 to the output of the feed-forward network block, giving the layer’s output representations *E_l_*. The whole process is repeated for each layer of the network. For notational simplicity, we also omitted the bias terms in our notations, but all linear operators except the initial embedding and positional encoding use a bias term. We refer to Vaswani *et al.* (2017) for further details of attention mechanisms and to Elnaggar *et al.* (2021) for details of the ProtBERT model.

#### 2.1.2 Decoder model

Transformer decoders use similar input embedding and positional encoding layers as the encoder, but their inputs are now the labels.

##### Initial Embedding

In the first step, decoder processes the output labels **y** = (*y*_1_,…, *y_n_*) similarly as the encoder processes the residues in the sequence **a** = (*a*_1_, …, *a_n_*). An important difference is that we append *y*_0_ = {BOS} (beginning of sequence) token, which will be used by the model to predict the first label *y*_1_. The matrix *Y* ∈ *R*^9×(*N* +1)^ consisting of the one-hot label representations as columns is mapped to a dense representation using *I_D_* = *W*_y_*Y* ∈ *R*^*d*×(*N*+1)^, where *W*_y_ ∈ *R*^*d*×9^ is the initial label embedding layer for all nine unique labels (eight real labels and {BOS}).

##### Positional encoding

We use a different type of positional encoding for the decoder part of our model. One alternative to modeling positional vectors using a linear layer, as in Section 2.1.1, is to use a sinusoidal function. The resulting fixed matrix *W_S_* ∈ *R*^*d*_*s*_ ×(*N* +1)^ (for a sequence of length *N* with an additional element *y*_0_ = {BOS}) is defined element-wise as:

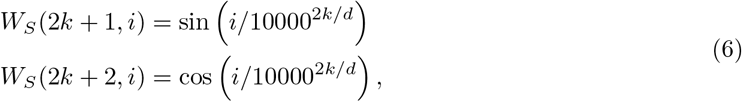

where 2*k* + 1 and 2*k* + 2 refer to the odd and even dimensions in our *d_S_*-dimensional positional encoding vector (here, *d_S_* = *d*) with *k* ∈ {0,…, *d_S_*/2 – 1} and *i* ∈ {1,…, *N* +1} is the sequence residue index.

For the decoder, the fixed *d_S_*-dimensional positional information *W_S_* is concatenated with the *d*-dimensional vector representations of the labels *I_D_*. The initial label sequence representation matrix is given by *D*_0_ = *I_D_* ⊕ *W_S_* ∈ *R*^(*d*+*ds*)×(*N* +1)^, where ⊕ denotes the concatenation operator and the sequence length is increased by one to *N* + 1 because of the additional *y*_0_ token. We choose this sinusoidal positional encoding for the decoder as it does not need any further training. Because of the limited amount of data, we hypothesize that another linear positional encoding would be difficult to converge well (note that the linear positional encoding in the encoder is already pre-trained on large amounts of data).

We concatenate an additional d-dimensional vector **g** ∈ *R*^*d*×1^ representing the organism group of the input sequence (obtained using a linear layer *W_G_* ∈ *R*^*d*×4^) to the encoder’s final layer residue representations *E_L_*. The result is also concatenated with the same sinusoidal positional encoding *W_S_*, in order to have the same type of positional information in the label and residue representations, forming 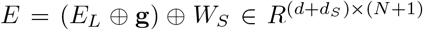, where the first concatenation is along the sequence dimension *N* for *E_L_* ∈ *R^d×N^* and **g**^*T*^ ∈ *R*^*d*×1^ and the second one along the model’s dimension d (see Supplementary Section 3 for the predictive performance effect of the additional positional encoding used on the encoder’s outputs).

##### Transformer decoder

Compared to the encoder layers, the decoder contains both a self-attention module, as well as an additional cross-attention layer which “looks” at the input vectors from the encoder. The first step of a decoder layer is a self multi-head attention, similar to the transformer encoder. Label representations are contextualized using the matrix from the previous layer *D*_*l*-1_, as described in Equations 4–5 (with *E_l_* replaced by *D_l_*). We denote this intermediate matrix of the decoder as 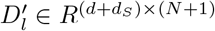. The label vector representations of 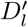 are then used as queries in the second multi-head attention layer, together with the representations from the encoder *E* ∈ *R*^(*d*+*d_s_*)×(*N* + 1)^. Concretely, similarly as in Equation 4, values from 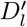 mapped into queries are attending to key and value mappings from *E* ∈ *R*^(*d*+*d_s_*)×(*N*+1)^ to yield an additional intermediate representation 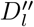. Lastly, a two-layered feed-forward network retrieves the next layer’s representations of the decoder *D_l_* ∈ *R*^(*d*+*d_s_*)×(*N* +1)^, like the ones in the encoder layers.

During training, the self attention module in the decoder layers computes all contextual values at once (all pairs (*y_i_*,*y_j_*) are considered) and therefore vectors representing *y_i_* have access to *y_j_,j* ≥ *i*. This induces undesired behaviour, as we wish to extract the amino acid labels based only on the previous labels. This issue is addressed by adding an additional {BOS} (beginning of sequence) token at the start of the label sequences and using appropriate masking. The mask is a matrix filled with zeros and –∞ above the diagonal and it can be added to the unnormalized attention weights 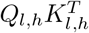 (see Equation 5). This ensures that the attention weights of current and future labels will be 0, and *y*_1_ will be predicted based only on the input retrieved by the encoder (as {BOS} does not contain any label information). Padding also uses a similar masking approach, where the attention to {PAD} tokens are zeroed.

The type of SP can only be correctly predicted when considering all residues forming an SP, and this requires the model to capture long range context. In contrast, the exact cleavage site prediction can be hindered by the fact that close residues have similar representations because of the context, and therefore we concatenate a one-hot representation of residue *k* for the associated label prediction *ŷ_k_*. We hypothesize that this allows the model more freedom in terms of the optimal amount of context it can add in its representations, and we observed consistent improvements when using this addition, which supports this claim.

Label predictions *y_k_* are finally computed based on the encoder’s last layer representations (**e**_*L*,1_, **e**_*L*,2_, …, **e**_*L,N*_), the previously predicted labels during inference (and previous true labels during training) (*y*_0_,*ŷ*_1_,…, *ŷ*_*k*-1_), as well as an additional one-hot encoded residue **c**^(*k*)^ at the position where label k is being predicted. During inference, the model predicts *ŷ_k_* sequentially based on its own generated label sequence (*y*_0_,*ŷ*_1_,…, *ŷ*_*k*-1_) as well as the encoder representations

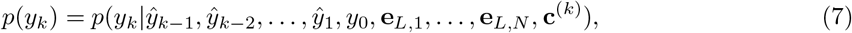

and *ŷ*_1_ will be predicted based the special token *y*_0_ = {BOS}, that we also use during training. Concretely, 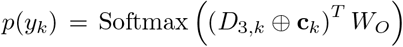, where *D*_3,*k*_ ∈ *R*^*d*+*d_s_* ×1^ is the decoder’s last layer representation of the previous residue label (since the outputs are shifted to the right by one position due to the {BOS} token), **c**_*k*_ ∈ *R*^21×1^ is the one-hot representation of the *k*^th^ amino acid and *W_O_* ∈ *R*^*d*+*d_s_*+21×8^ is a linear layer.

### 2.2 Architecture details

We use a dropout of 0.1 on all transformer decoder weights. The position-wise feed-forward network of our decoder has an almost fourfold expanding dimension, from the original 1152 to 4096 (*d* + *d_S_* = 1152, from the original representation *d* = 1024 and the concatenated sinusoidal positional information *d_S_* = 128). The ProtBERT encoder and transformer decoder have 30 and 3 layers of 16 attention heads, respectively. We initialize the decoder parameters with the Xavier Uniform initializer described by Glorot and Bengio (2010).

For better generalization, we used stochastic weight averaging which helps avoid sharp local minima solutions (Izmailov *et al.*, 2018). We chose constant learning rates of 10^-5^ and 10^-4^ for ProtBERT and decoder parameters respectively, ensuring as much exploration of the local minima as possible, without risking divergence. SWA weights are updated after each training epoch. For further details and insights on the effect of SWA we refer to Supplementary Section 4.

We also experimented with three other model variants that utilize the ProtBERT model in different ways. Based on the F1 scores from a CS prediction comparison, we chose the model setup presented in Section 2.1. See Supplementary Section 7 for details.

### 2.3 Dataset

We use the same dataset 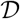 as Teufel *et al.* (2022), which contains sequences from Uniprot (Consortium, 2019) and Prosite (Sigrist *et al.*, 2012) for proteins containing SPs as well as UniProt and TOPDB (Dobson *et al.*, 2015) for soluble and transmembrane proteins, where only the expert-reviewed sequences are considered. The dataset contains 19174 protein sequences grouped into four organism groups: eukaryotes, gram-negative bacteria, gram-positive bacteria, and archaea. Every residue in each protein sequence has an annotated label that tells whether the residue belongs to the mature protein or the SP which will be cleaved, as well as the type of SP 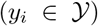. We use the same three-fold homology partitioning 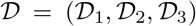, with the exception that we further split each partition into 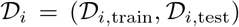, where 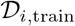 have 90% of the original 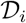 and 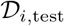 the re-maining 10%. We then train the model on 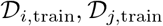, validate on 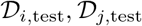, and then test on 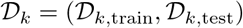. We therefore train and validate on different homology partitions than the test partition, ensuring a fair comparison against SignalP 6.0.

We do not use all the sequences in 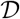 for the benchmark comparisons. Instead, we compare the predictive performance of TSignal and all other models on a benchmark subset 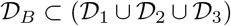. 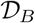 was created in SignalP 5 (Almagro Armenteros *et al.*, 2019) such that sequences in 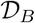 have at most 25% sequence identity to the training data set used by DeepSig (Savojardo *et al.*, 2018), and therefore all comparisons between TSignal, DeepSig and SignalP 6.0 are fair.

### 2.4 Evaluation

For cleavage site prediction, we use precision and recall as our main performance evaluation metrics. The precision and recall are computed for each SP type and organism group individually. We consider a cleavage site prediction to be correct if *i_p_* ∈ [*i_c_* — tol, *i_c_* + tol], where *i_p_* is the predicted index of the cleavage site, *i_c_* is the true (annotated) index and the tolerance tol ∈ {0,1, 2, 3}. Additionally, a cleavage site is only considered correct if the predicted signal peptide type is correct. For example, in the case where a signal peptide exists in a sequence but the predicted signal peptide type is not correct, the CS prediction is accounted as both a false positive (for the SP type which is wrongly predicted) and a false negative (for the true SP type). To have a single metric for the model’s performance, we use F1 score defined as 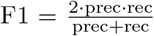. To summarize the results across all signal peptide types and organism groups, we report the average F1 score and weighted F1 score (weighted by the number of data points in each group).

To assess the SP presence prediction performance, we use Matthew’s correlation coefficient (MCC). We compute two separate metrics, MCC1 which considers only soluble and transmembrane proteins as negative samples, and MCC2 which also counts other SP types as negative samples.

## 3 Results

In Sections 3.1 and 3.2 we report benchmark comparisons on Sec/SPase I and II and Tat/SPase I sequences (because only these SP types have numeric results reported in (Teufel *et al.*, 2022)) from 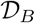 described in Section 2.3. This allows us to directly compare our results to the results of the previous state-of-the-art method, SignalP 6.0. The CS-F1 performance on the whole data 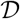 is reported in Supplementary Section 8, where we also report the Sec/SPase IV and Tat/SPase II performance.

### 3.1 Signal peptide prediction comparisons

We first assess the SP prediction performance on the sequences found in 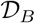 using the MCC metric. We evaluate MCC1 for TSignal, SignalP 6.0 and a few other popular models for SP prediction: DeepSig (Savojardo *et al.*, 2018), PRED-TAT (Bagos *et al.*, 2010), LipoP (Juncker *et al.*, 2003), and Phobius (Käll *et al.*, 2004). For the task of separating various types of SPs (MCC2), we only compare TSignal to SignalP 6.0 (because other models were not trained to distinguish all SP types considered in this work). Figure 3 shows the MCC values (we report the numeric values achieved by TSignal and SignalP 6.0 in Supplementary Table 4). We observe small but consistent improvements on Sec and Tat SPase I, and a slight decrease in Sec/Spase II SP type prediction performance compared to most of the previous approaches.As we show later, the model learns to detect the RR motif for Tat predictions, so the increased Tat/SPase I performance suggest our model finds causal features, important for good generalization. The weighted MCC1 and MCC2 scores across all organism groups and SP types for TSignal are 0.8579 ± 0.0022 and 0.8396 ± 0.014 respectively, while SignalP 6.0 has 0.8578 and 0.8365. We therefore note similar, or even a slight improvement, on the SP type prediction accuracy compared to previous state-of-the-art method.

**Figure 3:**
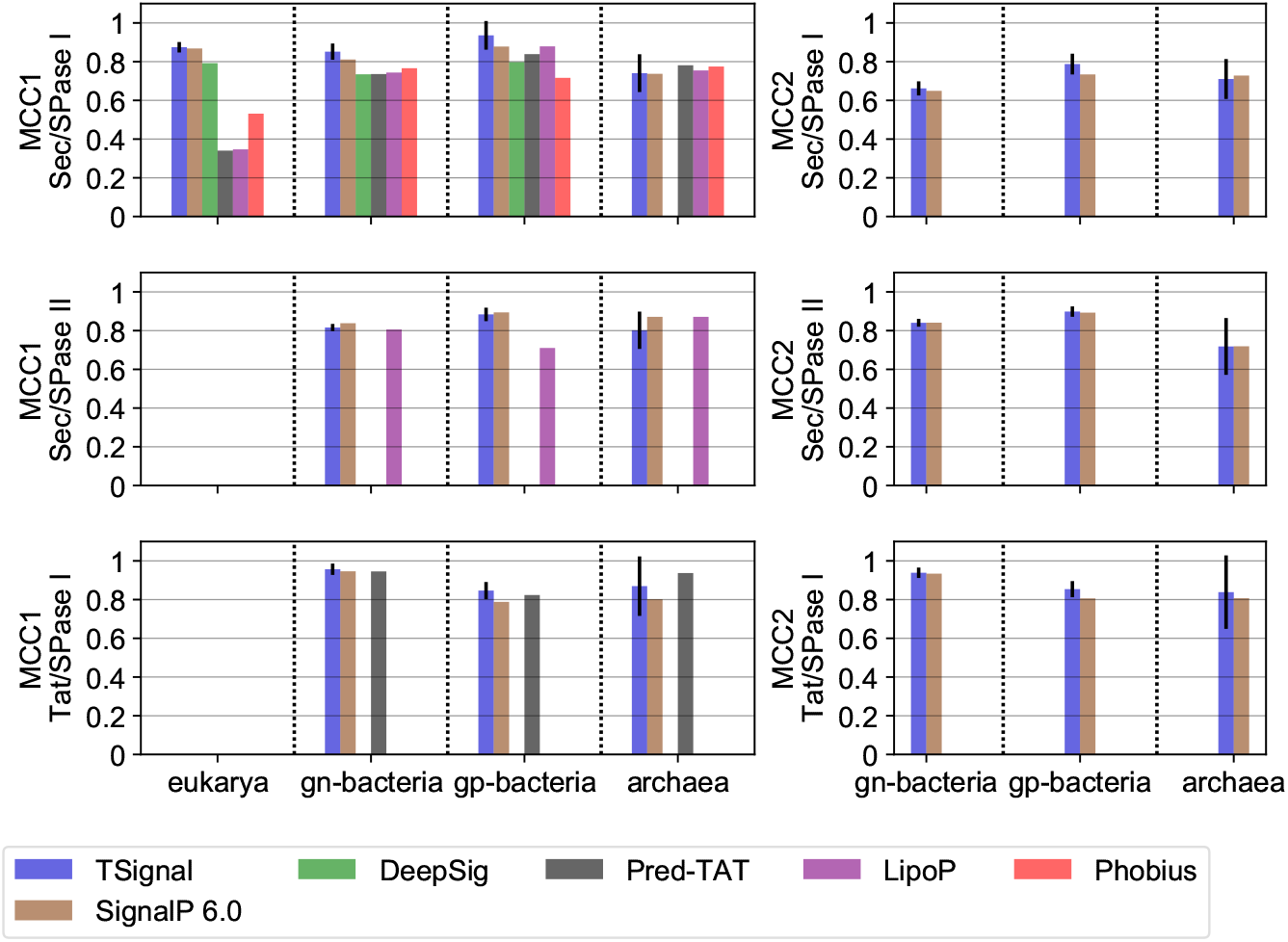
A comparison of SP predictions for TSignal and other models on the benchmark dataset 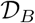 using the average MCC metrics: MCC1 on left and, MCC2 on right. The height of the bar plots represent the mean MCC result across five different runs, and the approximated 95% confidence interval is shown by the black vertical lines plotted on top of the bars. Organism groups are shown on the x-axis, and the SP-type that is tested on the y-axis. The benchmark data set 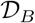 is homology partitioned only between the train and test set of TSignal, SignalP 6.0 and DeepSig, and results for other models are likely overestimates. Missing bars in the plots correspond to the respective model not being trained on that particular organism group or SP type.

To further test TSignal’s ability to recognize difficult-to-predict signal peptides, we assessed its capability in identifying four signal peptide-containing sequences that were earlier identified in a separate study that will be published separately (Kellosalo *et al.* unpublished; see Supplementary Section 2 and Fig. 1). These four sequences were identified in a screen for functional signal peptides that mediate protein secretion in mammalian cells and were not recognized to contain an SP by existing prediction methods. Distinctively, all of these sequences contain basic amino acids dispersed throughout the signal peptide sequence. Basic residues are typically contained at the N terminus of SPs, yet these previously unidentified sequences are sufficient to facilitate protein secretion in human cells (Supplementary Fig. 2). We postulate that in this case imposing specific transitions through structured prediction models may hinder the prediction of unusual sequences. Here, we tested the same models we compare against for the SP type and CS predictive performance using their publicly available webservers: DeepSig, PRED-TAT, LipoP, Phobius, and SignalP 6.0. DeepSig classifies three of these sequences as containing a transmembrane (TM) domain and Phobius reports a TM domain in all four sequences, but none of the methods classifies any of those four sequences as containing a SP. By contrast, TSignal predicts a TM domain in two of the four sequences and correctly determines a SP in the other two. The four sequences and their TSignal predictions are reported in Supplementary Fig. 2.

### 3.2 Cleavage site prediction comparison

We now test the CS prediction accuracy using the CS-F1 score. We only compare against SignalP 6.0 as the other methods do not predict all three SP types considered. Figure 4 shows that our model compares favourably to SignalP 6.0 for most organism groups and SP types on 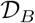 (we report the numeric values of the F1 score, precision and recall in Supplementary Tables 1, 2 and 3). Particularly interesting is that the cleavage site of Sec/SPase II SPs are more accurately predicted by our fully data-driven model, although the presence of Cys residues restricts the number of possible cleavage sites, and should help structured prediction models. We additionally note that Tat CS predictions are also better, even though the CRF model SignalP 6.0 is explicitly trained to detect the twin-arginine motif. In terms of overall performance, TSignal outperforms SignalP 6.0 on the majority of signal peptide types and organism groups with weighted F1 score of 0.8127 ± 0.005 compared to SignalP 6.0 0.7976.

**Figure 4:**
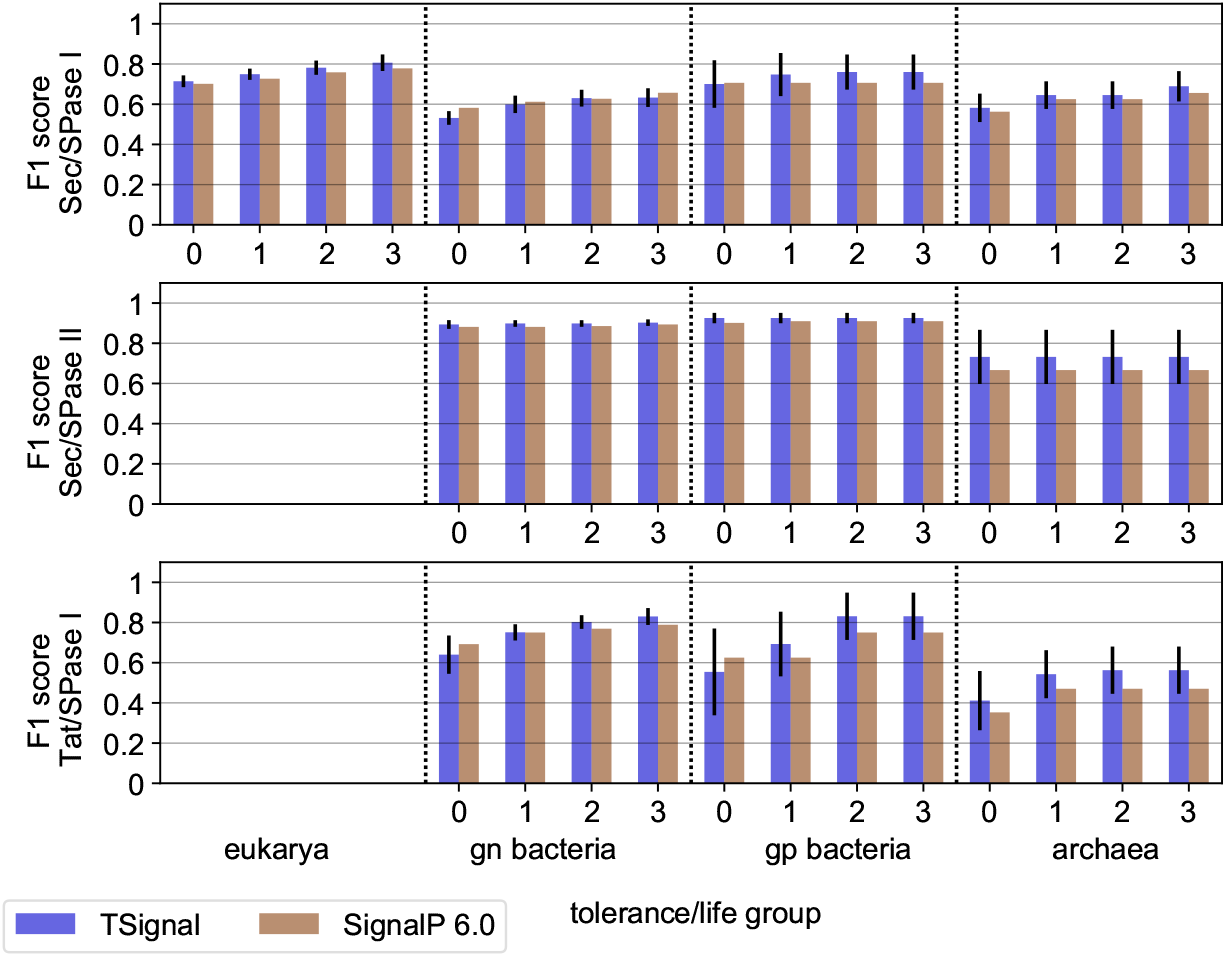
A comparison of CS predictions between TSignal and SignalP 6.0 on the benchmark dataset 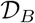 using the average F1 score. The height of the bar plots represent the mean F1 result across 5 different runs, and the approximated 95% confidence interval is shown by the black vertical lines plotted on top of the bars. Organism groups and tolerance levels are shown on the x-axis, and the SP-type that is tested on the y-axis. SignalP 6.0 results were computed using the precision and recall scores reported in their manuscript.

We also compare TSignal to PRED-TAT, LipoP, Phobius, and DeepSig on Sec/SPase I and no-SP sequences from 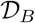. The results for these models were assessed using their publicly available web servers. We only use Sec/SPase I sequences since all these models have been trained (at least) on Sec/SPase I and no-SP sequences. We report these results in Figure 5. Note that 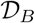 was homology partitioned to DeepSig’s training data in (Almagro Armenteros *et al.*, 2019), so comparing TSignal to it is fair, while all other results may be overestimates, due to the lack of homology-based test set partitioning of this experiment.

**Figure 5:**
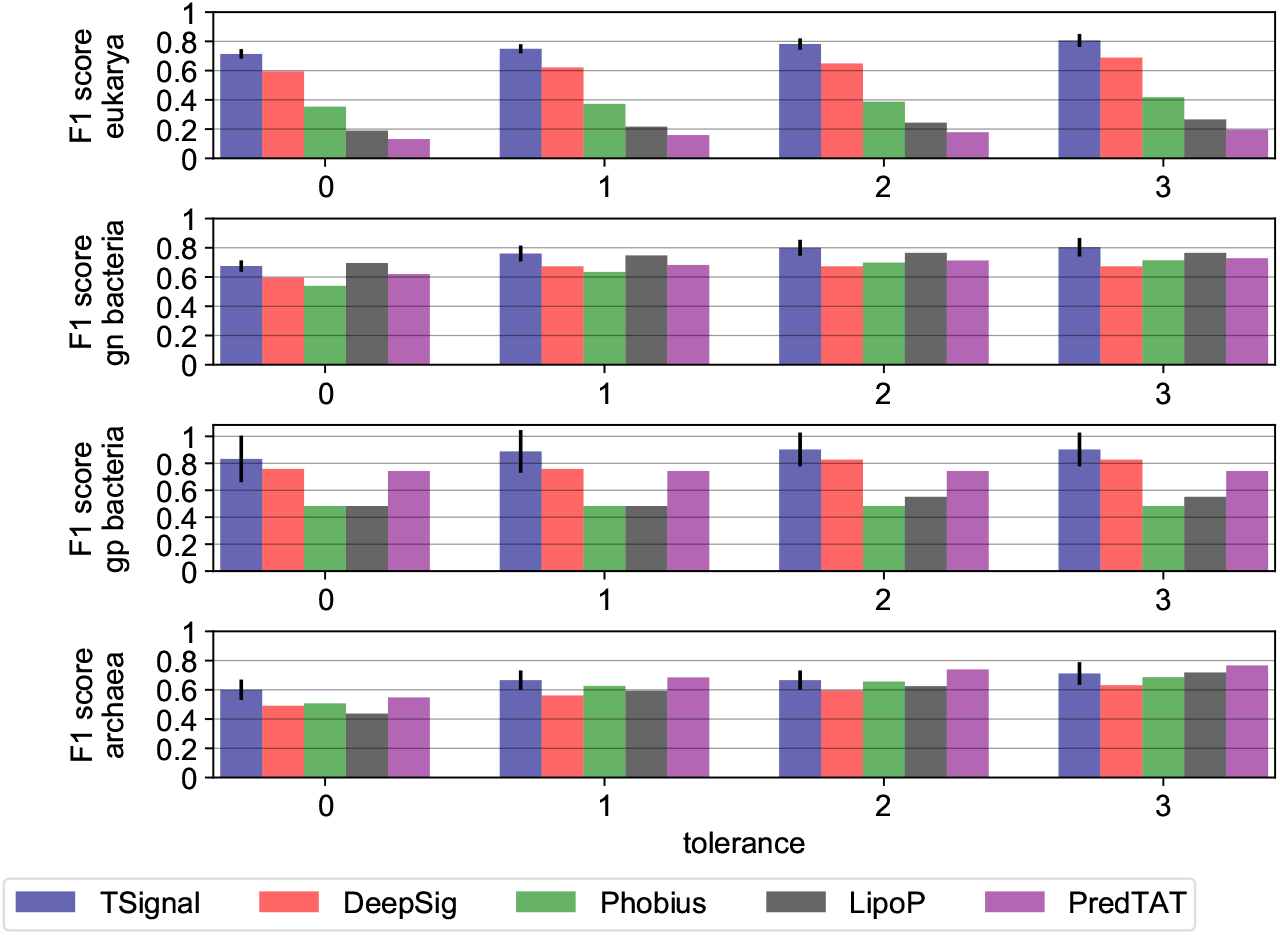
A comparison of CS predictions between TSignal and other popular models. We use the publicly available website tools for each of the tested models. The height of the bar plots represent the mean F1 result across five different runs, and the approximated 95% confidence interval is shown by the black vertical lines plotted on top of the bars. Tolerance levels and organism groups are shown on the *x*-axis and *y*-axis, respectively. The results were computed using only Sec/SPI sequences from the benchmark dataset 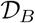, since the other models were not trained for all SP types considered here.

The predictions of TSignal are carried on diverse sequences, as we predict the sequences on the homology split test set, and from those extract the no-SP and Sec/SPase I sequences present in 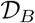. Although the models we compare against in Figure 5 are likely trained on sequences that are similar to those we use to test them, we still notice considerable improvements across most tolerance levels for Sec/SPase I CS predictions, and therefore we can be fairly confident our model outperforms these previous methods. Note that we could not include SignalP 6.0 here, as that would require access to their predicted test sequence labels, which we do not have.

### 3.3 Model performance analysis

To assess the ability of TSignal model to learn and generalize useful and interpretable information about an SP when predicting its type and CS, we employ a similar approach as Simonyan *et al.* (2014). As our training procedure is fully data-driven, we investigate the model’s ability to learn the information which can be useful for structured prediction models. We compute the average input importance scores for each residue in aligned sequences. We investigate 26 Tat/SPase I sequences that have the “RRXFLK” motif and 1682 Sec/SPase II sequences. We align the Tat/SPase I sequences wrt. the RR motif and the Sec/SPase II wrt. the Cys residue that is always present in the first position after the CS. We denote *i*_*c*+1_ to be the first residue after the cleavage site. We compute the input importance scores of the predictions for the test set, to investigate whether this information is generalized on sequences that are dissimilar to those used in the training. We refer to Supplementary Section 6 for more details on how we extract residue importance scores.

In Figure 6, the top panel shows how the model distinguishes the twin-arginine motif. The twin-arginine motif clearly has a high relative importance in the Tat prediction. The bottom panel illustrates the relevance of the cysteine residue (positioned at *i*_*c*+1_) for Sec/SPase II CS predictions. We also align Sec/SPase I sequences on the CS and plot them together. The red curve (Sec/SPase I) shows no relative importance of the *i*_*c*+1_ residue compared to the other residues surrounding the CS, whereas in Sec/SPII, there is a very clear spike on the position matching the *i*_*c*+1_ (Cys residue).

**Figure 6:**
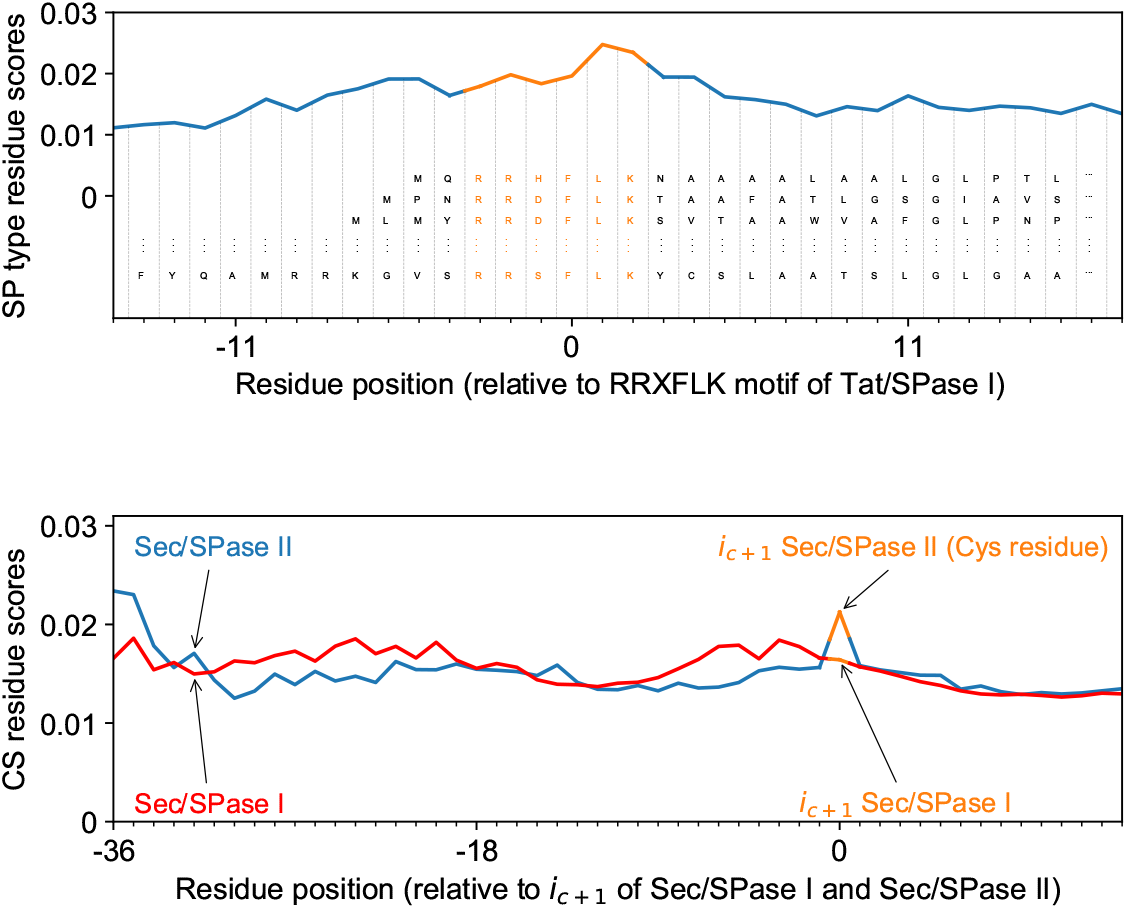
Input importance scores for (top) SP type predictions for Tat/SPase I and (bottom) CS predictions for Sec/SPase I and Sec/SPase II. On the top panel the sequences are aligned by the twin-arginine motif and it is shown how the model distinguishes the input embedding representation (denoted as *E*_0_ in this work) of the RR motif. On the bottom panel the sequences are aligned by the residue *i*_*c*+1_ following the CS. For Sec/SPase II residue *i*_*c*+1_ is cysteine and it has a high relative importance compared to the Sec/Spase I.

Next we evaluated how the performance of TSignal model increases with the amount of training data. Figure 7 shows that the model performance increases consistently as more training data is used, while the variance of the score estimates decreases. We assume that as more SP-protein pairs become available, the performance of our architecture will also steadily increase.

**Figure 7:**
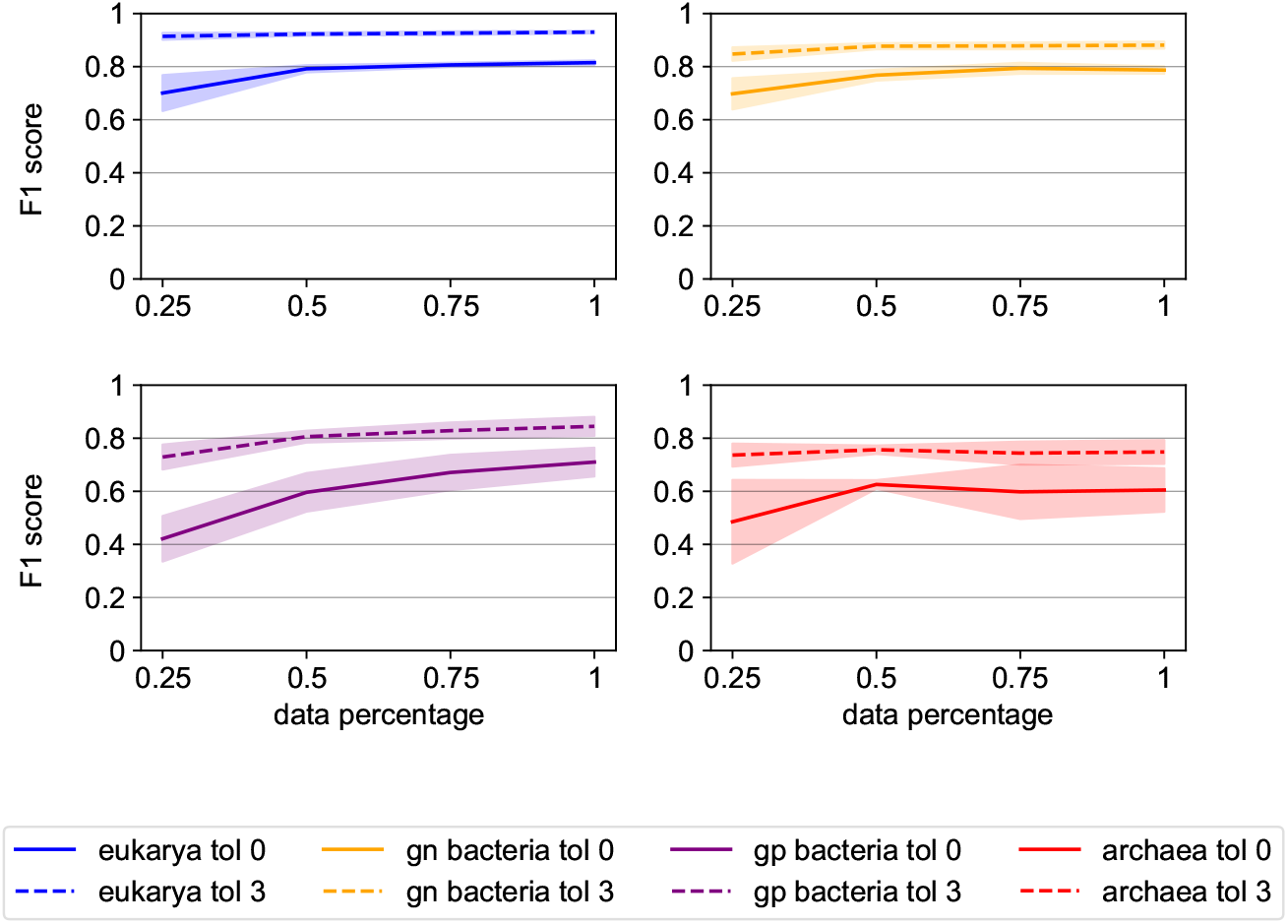
Performance of the TSignal model as a function of the amount of training data. F1 score is evaluated after training the model on 25, 50, 75 and 100% of the full training data while keeping the validation and testing data fixed. The test-train procedure is repeated five times, and we report the mean and 95% confidence interval (the shaded areas). Results are plotted for tolerances zero and three, for Sec/SPase I SPs from all organism groups.

We also check the model’s probability calibration. To a large degree the confidence scores of the model reflect the actual probability of the given prediction being correct in Supplementary Section 5.

## 4 Conclusion

We introduce, to our knowledge, the first deep learning model for signal peptide and cleavage site prediction, which does not use known biological properties of SPs explicitly (see Figure 1). Our results show that a transformer-based model provides competitive signal peptide prediction results and improves the accuracy of cleavage site prediction compared to the current state-of-the-art method. Indeed, on several organism groups, our transformer-based model outperforms previous methods. Our analysis also demonstrates that the model performance increases consistently with the amount of data. In other words, as more and more experimentally verified SP sequences will become available, data-driven end-to-end training of expressive deep learning models is likely to further improve the predictive performance. We also note that the amount of variability in TSignal’s performance is small, which indicates reliable performance evaluations as well as robust predictions.

Model interpretability is generally difficult to obtain for deep learning models and they are usually regarded as “black-boxes”. We show that our model generalizes biologically relevant information on homology partitioned data. In addition to the state-of-the-art cleavage site prediction performance, this further illustrates our model’s promising generalization potential on diverse sequences. Furthermore, it represents another argument for fully data-driven models, as information that was previously used in structured prediction models is learned by a model using a fully data-driven approach. We also note that hypotheses regarding the importance of various other motifs or specific residues could also be tested using the saliency map approach.

## Footnotes

1 We use *d* = 1024 but we use symbol *d* in method description for clarity.

2 This linear transformation is a very large input embedding layer that can encode positions in much larger sequences, but we preserve linear algebra notation for consistency.

