## Supplementary Material for "TSignal: A transformer model for signal peptide prediction"

### TSignal: A transformer model for signal peptide prediction
### Supplementary Material

Alexandru Dumitrescu, Emmi Jokinen, Juho Kellosalo
Ville Paavilainen, and Harri Lähdesmäki

#### 1 Result tables for TSignal and SignalP 6.0

Numeric CS and SP type performance metrics reported as tables. The numbers in these tables represent the mean and standard deviation results over 5 runs and are the same ones used in Figures 3 and 4 from the main text.

| | Sec/SPase I | | | | | | | | | | | | | | | |
|---|---|---|---|---|---|---|---|---|---|---|---|---|---|---|---|---|
| | eukarya | | | | gn-bacteria | | | | gp-bacteria | | | | archaea | | | |
| Tolerance | 0 | 1 | 2 | 3 | 0 | 1 | 2 | 3 | 0 | 1 | 2 | 3 | 0 | 1 | 2 | 3 |
| TSignal | **0.714 ± 0.010** | **0.749 ± 0.009** | **0.781 ± 0.013** | **0.806 ± 0.016** | 0.532 ± 0.012 | 0.599 ± 0.017 | **0.630 ± 0.016** | 0.632 ± 0.019 | 0.700 ± 0.055 | **0.748 ± 0.049** | **0.760 ± 0.039** | **0.760 ± 0.039** | **0.581 ± 0.031** | **0.645 ± 0.030** | **0.645 ± 0.030** | **0.689 ± 0.033** |
| SignalP 6.0 | 0.701 | 0.727 | 0.759 | 0.778 | **0.582** | **0.612** | 0.627 | **0.657** | **0.706** | 0.706 | 0.706 | 0.706 | 0.563 | 0.625 | 0.625 | 0.656 |
| | Sec/SPase II | | | | | | | | | | | | | | | |
| | eukarya | | | | gn-bacteria | | | | gp-bacteria | | | | archaea | | | |
| Tolerance | | | | | 0 | 1 | 2 | 3 | 0 | 1 | 2 | 3 | 0 | 1 | 2 | 3 |
| TSignal | | | | | **0.893 ± 0.006** | **0.898 ± 0.004** | **0.898 ± 0.004** | **0.902 ± 0.004** | **0.925 ± 0.008** | **0.925 ± 0.008** | **0.925 ± 0.008** | **0.925 ± 0.008** | **0.732 ± 0.062** | **0.732 ± 0.062** | **0.732 ± 0.062** | **0.732 ± 0.062** |
| SignalP 6.0 | | | | | 0.881 | 0.881 | 0.885 | 0.893 | 0.901 | 0.91 | 0.91 | 0.91 | 0.667 | 0.667 | 0.667 | 0.667 |
| | Tat/SPase I | | | | | | | | | | | | | | | |
| | eukarya | | | | gn-bacteria | | | | gp-bacteria | | | | archaea | | | |
| Tolerance | | | | | 0 | 1 | 2 | 3 | 0 | 1 | 2 | 3 | 0 | 1 | 2 | 3 |
| TSignal | | | | | 0.640 | **0.750 ± 0.016** | **0.802 ± 0.0126** | **0.830 ± 0.016** | 0.554 ± 0.103 | **0.693 ± 0.076** | **0.831 ± 0.0543** | **0.831 ± 0.054** | **0.411 ± 0.069** | **0.543** ± 0.056 | **0.563 ± 0.0543** | **0.563 ± 0.0543** |
| SignalP 6.0 | | | | | **0.692** | **0.75** | 0.769 | 0.789 | **0.625** | 0.625 | 0.75 | 0.75 | 0.353 | 0.47 | 0.47 | 0.47 |

Table 1: CS F1 scores for TSignal and SignalP 6.0 computed on the benchmark dataset $\mathcal{D}_B$.

| | Sec/SPase I | | | | | | | | | | | | | | | |
|---|---|---|---|---|---|---|---|---|---|---|---|---|---|---|---|---|
| | eukarya | | | | gn-bacteria | | | | gp-bacteria | | | | archaea | | | |
| Tolerance | 0 | 1 | 2 | 3 | 0 | 1 | 2 | 3 | 0 | 1 | 2 | 3 | 0 | 1 | 2 | 3 |
| TSignal | **0.68 ± 0.015** | **0.713 ± 0.015** | **0.744 ± 0.019** | **0.768 ± 0.022** | 0.466 ± 0.012 | 0.525 ± 0.018 | 0.552 ± 0.016 | 0.554 ± 0.017 | 0.624 ± 0.039 | **0.666 ± 0.037** | **0.678 ± 0.038** | **0.678 ± 0.038** | **0.678 ± 0.036** | **0.752 ± 0.04** | **0.752 ± 0.04** | **0.803 ± 0.034** |
| SignalP 6.0 | 0.661 | 0.685 | 0.715 | 0.733 | **0.534** | **0.562** | **0.575** | **0.603** | **0.632** | 0.632 | 0.632 | 0.632 | 0.643 | 0.714 | 0.714 | 0.75 |
| | Sec/SPase II | | | | | | | | | | | | | | | |
| | eukarya | | | | gn-bacteria | | | | gp-bacteria | | | | archaea | | | |
| Tolerance | | | | | 0 | 1 | 2 | 3 | 0 | 1 | 2 | 3 | 0 | 1 | 2 | 3 |
| TSignal | | | | | **0.952 ± 0.005** | **0.958 ± 0.003** | **0.958 ± 0.003** | **0.962 ± 0.003** | **0.976 ± 0.004** | **0.976 ± 0.004** | **0.976 ± 0.004** | **0.976 ± 0.004** | **0.788 ± 0.104** | **0.788 ± 0.104** | **0.788 ± 0.104** | **0.788 ± 0.104** |
| SignalP 6.0 | | | | | 0.913 | 0.913 | 0.917 | 0.925 | 0.929 | 0.938 | 0.938 | 0.938 | 0.583 | 0.583 | 0.583 | 0.583 |
| | Tat/SPase I | | | | | | | | | | | | | | | |
| | eukarya | | | | gn-bacteria | | | | gp-bacteria | | | | archaea | | | |
| Tolerance | | | | | 0 | 1 | 2 | 3 | 0 | 1 | 2 | 3 | 0 | 1 | 2 | 3 |
| TSignal | | | | | 0.645 ± 0.039 | **0.757 ± 0.017** | **0.809 ± 0.014** | **0.837 ± 0.014** | 0.641 ± 0.125 | **0.801 ± 0.101** | **0.96 ± 0.08** | **0.96 ± 0.08** | **0.452 ± 0.067** | **0.598 ± 0.057** | **0.618 ± 0.031** | **0.618 ± 0.031** |
| SignalP 6.0 | | | | | **0.679** | 0.736 | 0.755 | 0.774 | **0.714** | 0.714 | 0.857 | 0.857 | 0.375 | 0.5 | 0.5 | 0.5 |

Table 2: CS precision scores for TSignal and SignalP 6.0 computed on the benchmark dataset $\mathcal{D}_B$.

| | Sec/SPase I | | | | | | | | | | | | | | | |
|---|---|---|---|---|---|---|---|---|---|---|---|---|---|---|---|---|
| | eukarya | | | | gn-bacteria | | | | gp-bacteria | | | | archaea | | | |
| Tolerance | 0 | 1 | 2 | 3 | 0 | 1 | 2 | 3 | 0 | 1 | 2 | 3 | 0 | 1 | 2 | 3 |
| TSignal | **0.752 ± 0.005** | **0.789 ± 0.003** | **0.823 ± 0.007** | **0.849 ± 0.01** | 0.62 ± 0.016 | **0.698 ± 0.017** | **0.734 ± 0.019** | **0.738 ± 0.023** | **0.8 ± 0.084** | **0.853 ± 0.078** | **0.867 ± 0.06** | **0.867 ± 0.06** | **0.511 ± 0.038** | **0.567 ± 0.038** | **0.567 ± 0.038** | **0.606 ± 0.044** |
| SignalP 6.0 | 0.747 | 0.774 | 0.808 | 0.829 | **0.639** | 0.672 | 0.689 | 0.721 | **0.8** | 0.8 | 0.8 | 0.8 | 0.5 | 0.556 | 0.556 | 0.583 |
| | Sec/SPase II | | | | | | | | | | | | | | | |
| | eukarya | | | | gn-bacteria | | | | gp-bacteria | | | | archaea | | | |
| Tolerance | | | | | 0 | 1 | 2 | 3 | 0 | 1 | 2 | 3 | 0 | 1 | 2 | 3 |
| TSignal | | | | | 0.841 ± 0.008 | 0.846 ± 0.005 | 0.846 ± 0.005 | 0.85 ± 0.005 | **0.88 ± 0.015** | 0.88 ± 0.015 | 0.88 ± 0.015 | 0.88 ± 0.015 | 0.689 ± 0.044 | 0.689 ± 0.044 | 0.689 ± 0.044 | 0.689 ± 0.044 |
| SignalP 6.0 | | | | | **0.852** | **0.852** | **0.856** | **0.864** | 0.875 | **0.883** | **0.883** | **0.883** | **0.778** | **0.778** | **0.778** | **0.778** |
| | Tat/SPase I | | | | | | | | | | | | | | | |
| | eukarya | | | | gn-bacteria | | | | gp-bacteria | | | | archaea | | | |
| Tolerance | | | | | 0 | 1 | 2 | 3 | 0 | 1 | 2 | 3 | 0 | 1 | 2 | 3 |
| TSignal | | | | | 0.635 ± 0.047 | 0.745 ± 0.018 | **0.796 ± 0.016** | **0.824 ± 0.021** | 0.489 ± 0.089 | **0.611 ± 0.061** | **0.733 ± 0.042** | **0.733 ± 0.042** | **0.38 ± 0.075** | **0.5 ± 0.063** | **0.52 ± 0.075** | **0.52 ± 0.075** |
| SignalP 6.0 | | | | | **0.706** | **0.765** | 0.784 | 0.804 | **0.556** | 0.556 | 0.667 | 0.667 | 0.333 | 0.444 | 0.444 | 0.444 |

Table 3: CS recall scores for TSignal and SignalP 6.0 computed on the benchmark dataset $\mathcal{D}_B$.

| | Sec/SPase I | | | | | | |
|---|---|---|---|---|---|---|---|
| | eukarya | gn-bacteria | | gp-bacteria | | archaea | |
| | MCC1 | MCC1 | MCC2 | MCC1 | MCC2 | MCC1 | MCC2 |
| TSignal | $\mathbf{0.874 \pm 0.009}$ | $\mathbf{0.851 \pm 0.016}$ | $\mathbf{0.662 \pm 0.013}$ | $\mathbf{0.936 \pm 0.032}$ | $\mathbf{0.787 \pm 0.022}$ | $\mathbf{0.741 \pm 0.044}$ | $0.710 \pm 0.047$ |
| SignalP 6.0 | 0.868 | 0.811 | 0.649 | 0.878 | 0.734 | 0.737 | **0.728** |
| | | Sec/SPase II | | | | | |
| | | gn-bacteria | | gp-bacteria | | archaea | |
| | | MCC1 | MCC2 | MCC1 | MCC2 | MCC1 | MCC2 |
| TSignal | | $0.816 \pm 0.005$ | $0.840 \pm 0.006$ | $0.883 \pm 0.022$ | $\mathbf{0.898 \pm 0.009}$ | $0.802 \pm 0.044$ | $0.718 \pm 0.069$ |
| SignalP 6.0 | | **0.836** | **0.841** | **0.894** | 0.893 | **0.871** | **0.719** |
| | | Tat/SPase I | | | | | |
| | | gn-bacteria | | gp-bacteria | | archaea | |
| | | MCC1 | MCC2 | MCC1 | MCC2 | MCC1 | MCC2 |
| TSignal | | $\mathbf{0.957 \pm 0.010}$ | $\mathbf{0.939 \pm 0.009}$ | $\mathbf{0.846 \pm 0.018}$ | $\mathbf{0.854 \pm 0.016}$ | $\mathbf{0.869 \pm 0.072}$ | $\mathbf{0.839 \pm 0.090}$ |
| SignalP 6.0 | | 0.946 | 0.934 | 0.788 | 0.806 | 0.802 | 0.807 |

Table 4: SP type MCC1 and MCC2 scores scores for TSignal and SignalP 6.0 computed on the benchmark dataset $\mathcal{D}_B$.

### 2 Novel, experimentally verified eukaryote SPs

We tested five other popular SP prediction methods (SignalP 6.0, PRED-TAT, LipoP, Phobius, and DeepSig) on the four novel eukaryote SPs using their respective publicly available web servers. The tests were conducted with the corresponding available versions of these models from the 6th of May, 2022.

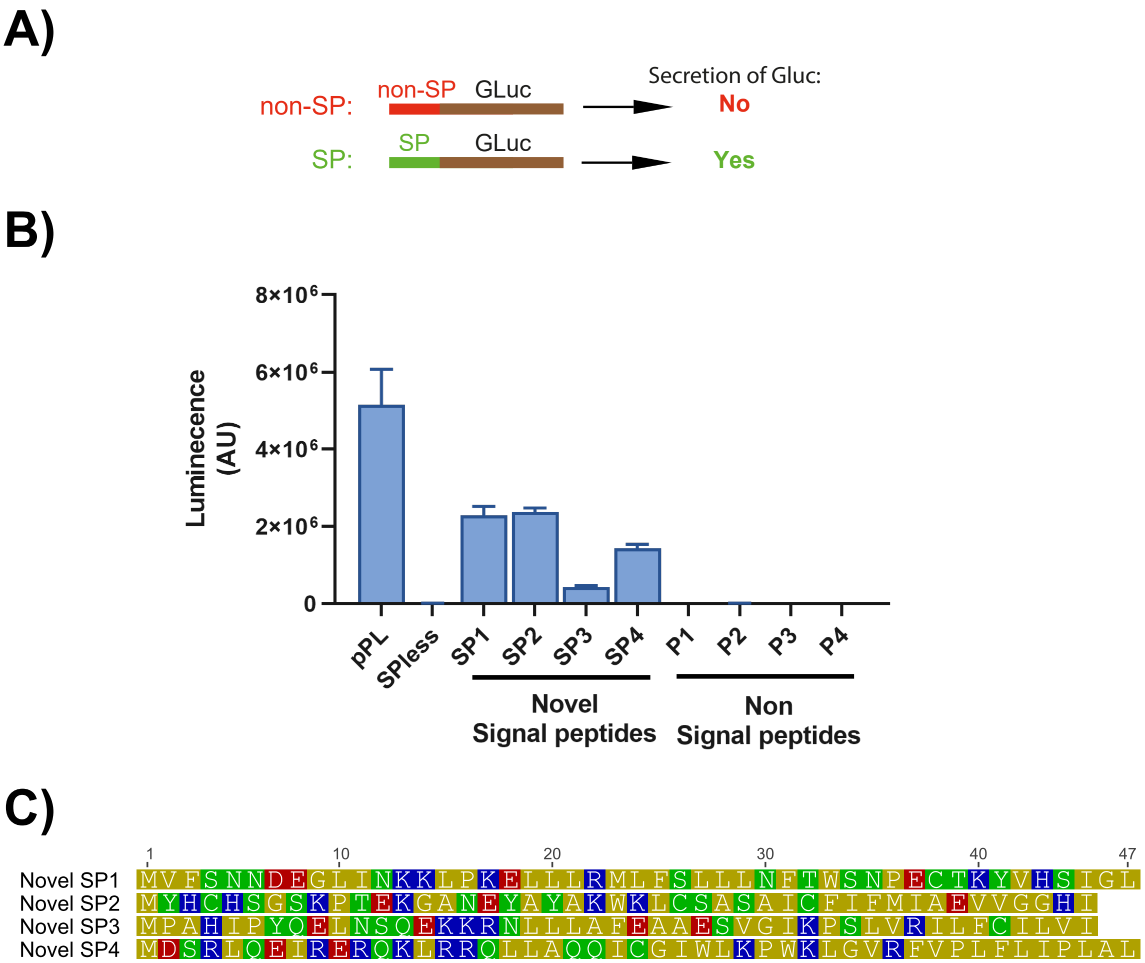


Figure 1: Functional signal peptide testing in human HEK293T cells. (A) schematic of the secreted Gaussia luciferase (GLuc) assay. GLuc constructs with appended N-terminal signal peptides are secreted into the cell culture medium, whereas constructs with no or non-signal peptide sequences are not secreted. (B) Luminescence of GLuc constructs containing different N-terminal peptides measured from cell media 26 hours after transfection. Full method description in section 5.2.1. (C) Primary sequences of tested, functional signal peptide-containing N-terminal peptide sequences

.

#### 2.1 Assessing signal-peptide function with secreted Gaussia luciferase

Different N-termini were fused to a Gaussia princeps luciferase (Gluc) that had been cloned into pcDNA5/FRT/TO. For this the Gluc-encoding plasmid was first digested with AgeI and NheI restriction enzymes (NEB). DNA constructs encoding a known non-SP sequence (MGTRSDST amino-acid sequence) or a known SP sequence (the human preprolactin SP with four additional mature-chain amino acids) were then cloned in to the linearized plasmid using a T4-ligase (NEB) (SPless and pPL-SP Gluc constructs, respectively), while the putative novel SP encoding DNAs were cloned in using the NEBuilder Hifi DNA Assembly (NEB). The sequences of all the cloned constructs were verified with Sanger sequencing.

For assessing if a cloned N-terminus contains a functional SP which can drive the secretion of Gluc,

```
    Input Sequence: MPAHIPYQELNSQEKKRNLLLAFEAAESVGIKPSLVRILFCILVITRATADAQSRMQIFVKTLTGKTITL
  Predicted Labels: OOOOOOOOOOOOOOOOOOOOOOOOOOOOOOOOOOOOOMMMMMMMMMMMMMMMMMIIIIIIIIIIIIIIII

    Input Sequence: MYHCHSGSKPTEKGANEYAYAKWKLCSASAICFIFMIAEVVGGHITRATADAQSRMQIFVKTLTGKTITL
  Predicted Labels: IIIIIIIIIIIIIIIIIIIIIIIIMMMMMMMMMMMMMMMMMMMMMOOOOOOOOOOOOOOOOOOOOOOOOO

    Input Sequence: MDSRLQEIRERQKLRRQLLAQQICGIWLKPWKLGVRFVPLFLIPLALTRATADAQSRMQIFVKTLTGKTI
  Predicted Labels: SSSSSSSSSSSSSSSSSSSSSSSSSSSSSSSSSSSSSSSSSSSSSSSSSSSSOOOOOOOOOOOOOOOOO

    Input Sequence: MVFSNNDEGLINKKLPKELLLRMLFSLLLNFTWSNPECTKYVHSIGLTRATADAQSRMQIFVKTLTGKTI
  Predicted Labels: SSSSSSSSSSSSSSSSSSSSSSSSSSSSSSSSSSOOOOOOOOOOOOOOOOOOOOOOOOOOOOOOOOOOOO
```

Figure 2: The resulted predictions for the four experimentally validated mammalian signal peptides obtained using TSignal.

we transfected HEK293T cells with the cloned plasmids. These transient transfections were done by first plating the cells on a 96-well plate by pipeting in to each well 100 µl cell suspension which contained $0.1 \cdot 10^6$ cells / ml in DMEM + 10% FBS (DMEM-FBS). The plated cells were then grown at 37° C under 5% CO2 for 24 hours after which the media was exchanged to fresh DMEM-FBS. Cells were then transfected with 150 ng DNA per well using 3.6 µl Transporter 5 (Polysciences) transfection reagent per well. Four separate wells were prepared for each transfected plasmid. 4h after transfection, the transfection reagent-containing media was removed, the cells were washed with 1xPBS and fresh DMEM-FBS was added on the cells. 22h after this media exchange, the cell media was collected and the luminescence of the secreted Gluc was measured with the Gaussia Glow-Juice Luciferase assay kit (p.j.k) on an Enspire multimode plate reader (Perkin Elmer).

#### 3 Additional positional encoding effect

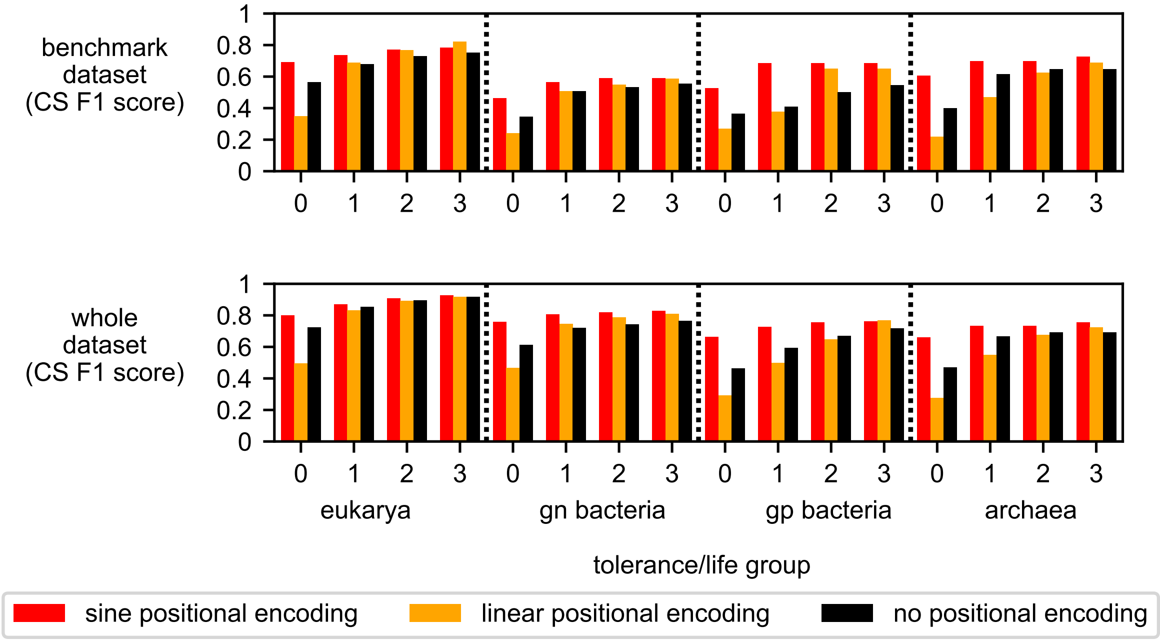


Figure 3: Analysis of performance increase when adding sinusoidal positional encoding to the residue vector representations (from ProtBERT) before using them as kay and value vectors in the decoder. The cleavage site F1 scores are computed for all tolerances and all organism groups for Sec/SPase I. We compare adding a sine-based extra positional encoding, an extra linear positional encoding and no extra positional encoding at all. These experiments were conducted with additive positional encodings.

#### 4 SWA effect

Stochastic weight averaging has been shown to improve generalization through finding wider local optima. The method uses the fact that under certain assumptions, weights retrieved by stochastic gradient methods are samples from the posterior probability $p(\mathbf{w}|\mathcal{D})$. Therefore, averaging the parameters $\mathbf{w}_i$ at multiple checkpoints $i$ during training should give a wider, less prone to overfitting solution. Compared to dropout, weight decay or other forms of regularization, SWA only requires selecting good learning rates, instead of e.g. extensive grid or random search for new hyperparameters.

To select an appropriate learning rate for the SWA procedure, we loaded and further trained an already converged model and determined the minimum learning rate that made the model diverge. The ProtBERT parameters were a lot more sensitive to an increased learning rate. The model diverges with an encoder learning rate above $2 \cdot 10^{-5}$, while the decoder learning rate can be increased up to $3 \cdot 10^{-4}$ without leaving the local optima. We choose learning rates of $10^{-5}$ and $10^{-4}$ for ProtBERT and decoder parts of TSignal respectively, ensuring as much exploration as possible without risking divergence.

We first train using early stopping based on the validation CS-F1 score, which stops the model at some epoch *ep*. We further tune the model from the checkpoint epoch *ep* using SWA for $ep/2$ more epochs.

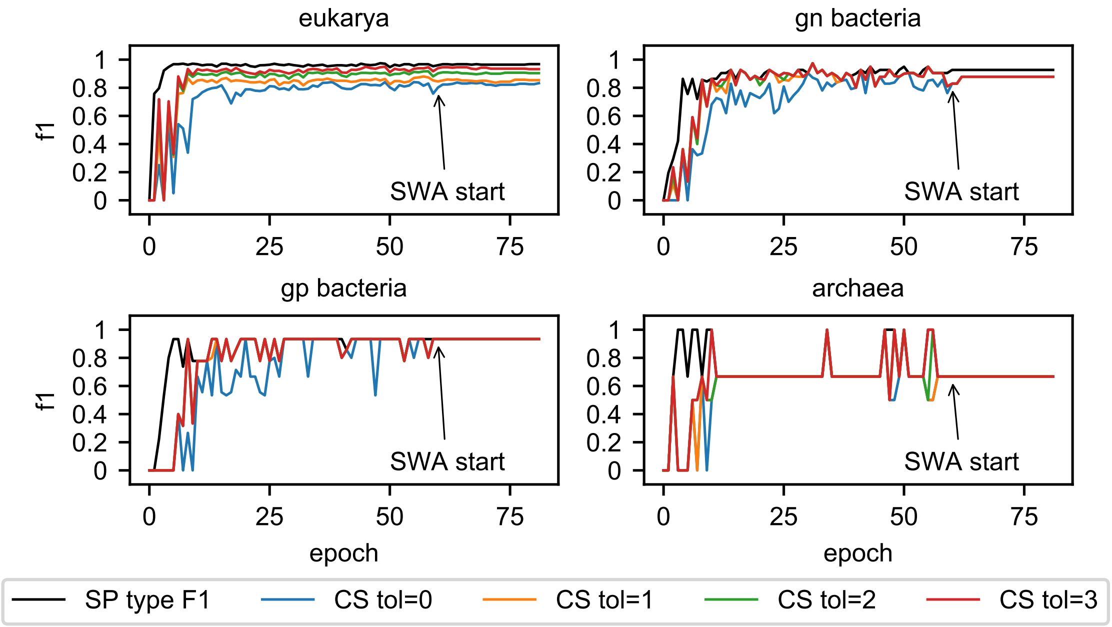


Figure 4: Validation F1 scores for Sec/SPase I over the epochs. SWA has a clear stabilization effect and helps the model generalize better through finding a better local mode.

### 5 Probability calibration

The prediction values being higher (more confident) in classification tasks, does not necessarily reflect a higher chance of the actual prediction being correct. Probability miscalibration is a common problem in very deep architectures. Although techniques like batch and layer normalization (Ioffe and Szegedy, 2015; Ba *et al.*, 2016) or skip connections proved useful in helping these large models converge faster and generalize better, it has been shown that performance increments achieved by more and more complex models come at the cost of not yielding useful confidence scores (Guo *et al.*, 2017).

In Equation 1, the interval $[0,1]$ (the possible values for the classification predictions) is divided into $M$ bins $B_m$. Then, $\mathrm{acc}(B_m)$ is the computed (empirical) accuracy of the model when its prediction probabilities are found in that interval $p \in B_m$. Similarly, $\mathrm{conf}(B_m)$ is the associated probability of bin $m$ (the mid point of the interval $B_m$). In a perfectly calibrated model, its confidence scores would be equal to the computed accuracy, and the resulting expected calibration error ECE would be 0. Visually, this means that the red bars in Figure 5 perfectly match the blue ones. We show the probability calibration of our model for the CS prediction, where we consider the CS probability to be the predicted probability $p(\hat{\mathbf{y}} \in \{\text{intracellular, extracellular, transmembrane}\})$, after a sequence of SP predictions. The ECE is defined as:

$$\mathrm{ECE} = \sum_{m=1}^{M} \frac{|B_m|}{n} \left|\mathrm{acc}\left(B_m\right) - \mathrm{conf}\left(B_m\right)\right|. \tag{1}$$

We estimated to what degree the probabilities of our model are miscalibrated. We evaluated ECE for the cleavage site prediction probability calibration for Sec/SPase I predictions for various tolerance levels. Figure 5 shows that the prediction results are approximately calibrated for different tolerance levels: TSignal is slightly overconfident for tolerance values 0 and 1, and nearly perfectly calibrated for tolerance values 2 and 3.

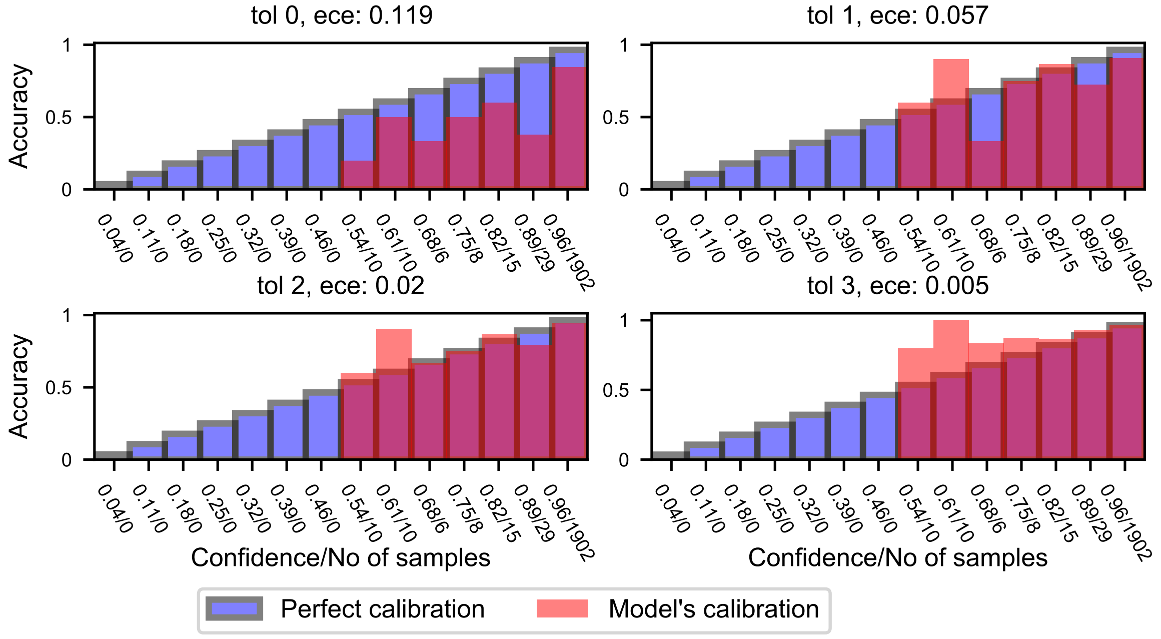


Figure 5: Cleavage site prediction probability calibration for Sec/SPase I predictions for various tolerance levels. Calibration error is assessed using the expected calibration error (ECE).

### 6 Residue saliency maps

We approximate the importance of a specific residue for a given prediction by taking the gradients of that prediction wrt. the input representations fed to the ProtBERT model, described in Equation 1 of the main text (and denoted as $E_0$). For both CS and SP type predictions we extract $\hat{\boldsymbol{y}}[i_{c+1}]$ and $\hat{\boldsymbol{y}}_1$, where $\hat{\boldsymbol{y}}[i_{c+1}]$ is the vector of predicted label probabilities of the first non-SP prediction $i_{c+1}$ (which we consider as a CS prediction) and $\hat{\boldsymbol{y}}_1$ is the first label prediction in a sequence, which determines the SP type.

We assess the average importance over multiple sequences, and therefore align the sequences wrt. the true CS when determining the cysteine importance in Sec/SPase II CS predictions $\hat{\boldsymbol{y}}[i_{c+1}]$. Similarly, we align Tat directed sequences according to the "RRXFLK" motif when checking the residue importance of the SP type prediction given by the gradients of $\hat{\boldsymbol{y}}_1$. A visual illustration for the alignment for Tat sequences is shown in Figure 6. The equation describing the average importance scores for residues $r_i$ at position $i$ (relative to CS or the RR motif) over multiple sequences is given below:

$$\mathrm{IS}_{r_i} = \frac{1}{N_i}\frac{1}{d}\sum_{j=1}^{N_i}\sum_{k=1}^{d} \mathrm{abs}\left[\nabla_{E_{0,i}^{j,k}}\left(\max_{c\in\mathcal{Y}} p(\hat{\boldsymbol{y}}_{t,j}=c)\right)\right], \tag{2}$$

where $N_i$ is the total number of residues at position $i$ across the tested sequences, $d$ is the model's dimension, $E_{0,i}^{j,k}$ is the input residue representation $E_0$ of residues at the (relative) position $i$, in the $j^{\text{th}}$ sequence, on the $k^{\text{th}}$ dimension of our $d$-dimensional input vector representation of residues, and $p(\hat{\boldsymbol{y}}_{t,j}=c)$ is the model's predicted probability that the first residue (SP type prediction, $t=1$) or the CS residue ($t=i_{c+1}$) of sequence $j$ belongs to class $c$.

In Figure 6 we give a simplified example of three Tat sequences aligned to the RR motif, which shows $N_i$ being computed separately for each relative position $i$. Note that this is crucially different than simply dividing the summed importance scores (gradients) of the sequence residues to the total number of sequences. A similar approach was employed for the CS alignment in the Sec/SPase II analysis we made.

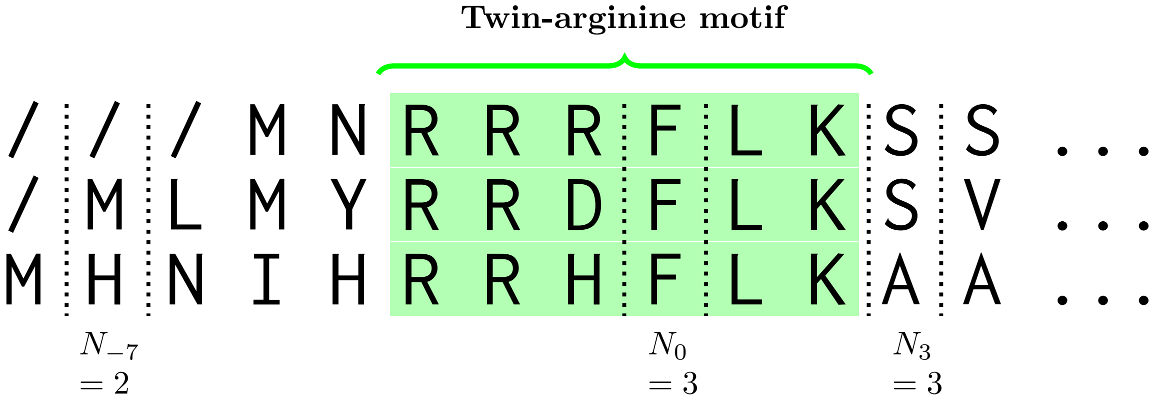


Figure 6: Illustration of the Twin-arginine alignment method. $N_0$ is arbitrarily chosen as the "center" of the RR motif at the phenylalanine residue, and all other indices are relative to $N_0$. The length of the preceding and following residues relative to the center of the motif varies, so the average importance scores have to be taken considering the number of residues $N_i$ at each relative position $i$.

### 7 Different model variants

We use the ProtBERT model of Elnaggar *et al.* (2021) and experiment with four transformer model variations. The best performing approach is using the ProtBERT LM as the encoder of a modified encoder-decoder architecture from Vaswani *et al.* (2017). The ProtBERT residue representations are used in the second multi-head attention of each of the decoder's layers as keys and values, as described in the main text. Masked token prediction (and next sentence prediction in natural language) enabled BERT LMs to encode rich representations of tokens using unsupervised learning. Note however that BERT models use an identical architecture as the encoder of Vaswani *et al.* (2017). Therefore, the method we present uses a form of transfer learning, where the transformer encoder is initialized with the pre-trained ProtBERT weights. This is a crucial aspect that enables these highly complex transformer architectures to be trained on the relatively small SP dataset we use.

In the first experiment, we consider a randomly initialized 3-layered encoder-decoder transformer architecture. The inputs of this architecture are retrieved by ProtBERT, but we do not further fine-tune the LM's weights. This approach would be equivalent to replacing the input embeddings denoted as $E_0$ in the main text with ProtBERT's residue representations (which are fixed in this approach, as ProtBERT does not change during training).

In the second experiment we tune the ProtBERT weights on the SP-CS prediction task separately, and then extract the tuned representations of the resulting protein LM to train a 3-layered encoder-decoder, similar to the first experiment. To fine-tune ProtBERT (separately), we use the fact that the input and output sequences have the same length - we extract the ProtBERT LM's representations from the last layer, having $N$ vectors summarized by $E_{30} \in R^{N \times d}$. Each vector is passed to a linear layer that predicts a corresponding label $\hat{y}_k$, and the gradients w.r.t. the cross entropy loss between $\hat{y}_k$ and the true $y_k$ flow through the ProtBERT parameters.

We then experiment with tuning ProtBERT along with our randomly initialized encoder-decoder architecture, effectively extending the number of our encoder's layers to 33 (ProtBERT having 30 layers to which three additional layers are added from the randomly initialized encoder-decoder architecture).

Removing the additional encoder layers yields the final model, which achieved the best results. We tune ProtBERT and the transformer decoder layers together, using the same Adam optimizer, but with separate learning rates.

Additionally, our experiments showed that when we do not tune the ProtBERT model (either prior tuning or training ProtBERT together with TSignal), the results are better if we separate the SP type predictions from the sequence predictions (experiments a and b). We do this by training TSignal with an agnostic "S" label, for any signal peptide, and then have a second convolutional model predict the signal peptide type. The architecture used for the signal peptide type prediction is similar to the convolutional architecture of Gligorijević *et al.* (2021). We report the comparative CS-F1 performance results in Figure 8.

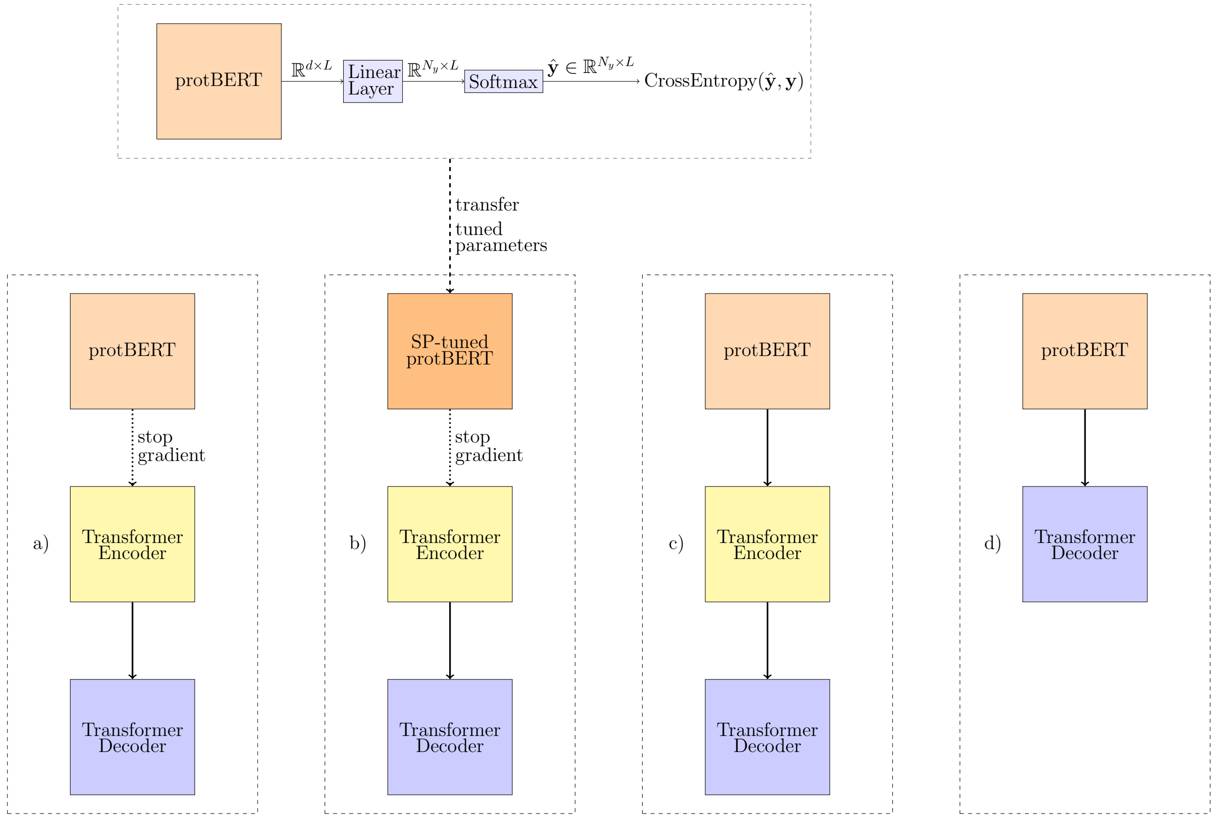


Figure 7: The four types of models that we tested. a) ProtBERT model is not further tuned, and its parameters are not modified during the 3-layered encoder-decoder model training. b) ProtBERT parameters are separately tuned on the SP label prediction task and an encoder-decoder transformer architecture is trained using the tuned ProtBERT representations as inputs. c) ProtBERT is part of the model, trained together with the rest of the parameters. d) ProtBERT is treated as the encoder of the transformer encoder-decoder architecture, with no other additional encoder parameters being added.

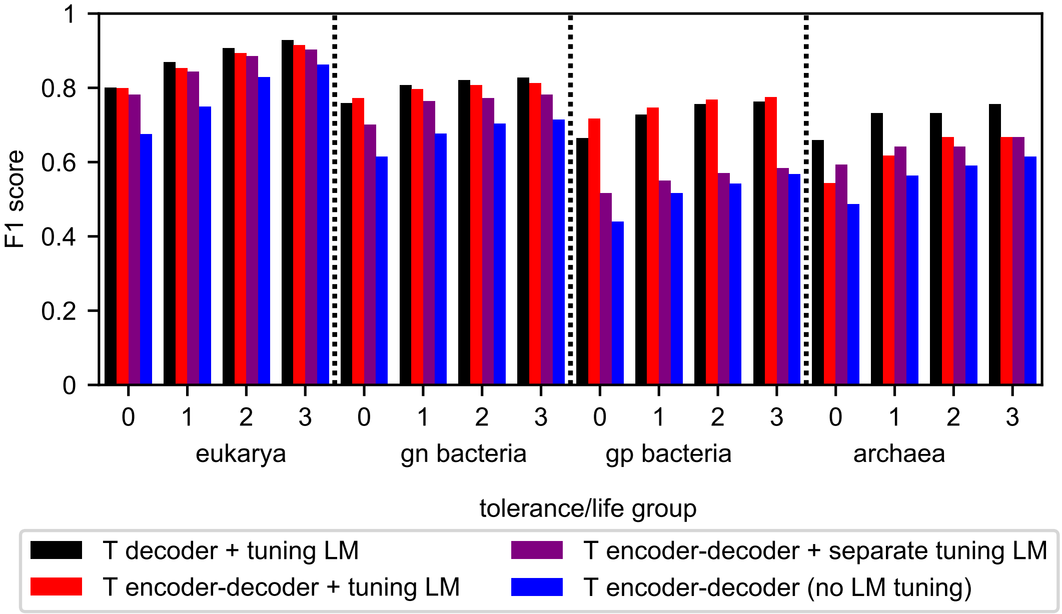


Figure 8: Comparison between the experimented methods. The best performance is achieved when ProtBERT is used as the encoder of our transformer model. We also test adding a three additional layers to the ProtBERT LM, effectively creating a 33-layered transformer encoder that is used by our 3-layered decoder. Tuning ProtBERT separately, yielded worse results, but it still performs better than using only the raw ProtBERT embeddings, without modifying its weights at all.

### 8 Performance on all data from $\mathcal{D}$

We measured the model's F1 score on the whole homology partitioned data to retrieve a reliable performance estimation for our model. We ran the model training five times and plotted the mean and 95% confidence interval in the shaded area in Figures 9, 10, and 11. The final test performance is very stable, having a low test performance variance, especially for organism groups and SP types where there is a reasonably high amount of training sequences. These results and the ones related to the performance score as a function of training data amount, show that our model should have better and more reliable performance as more data becomes available in the future. We could not compare this against SignalP 6.0, as they do not report numeric results for Tat/SPase II and Sec/SPase IV SP types.

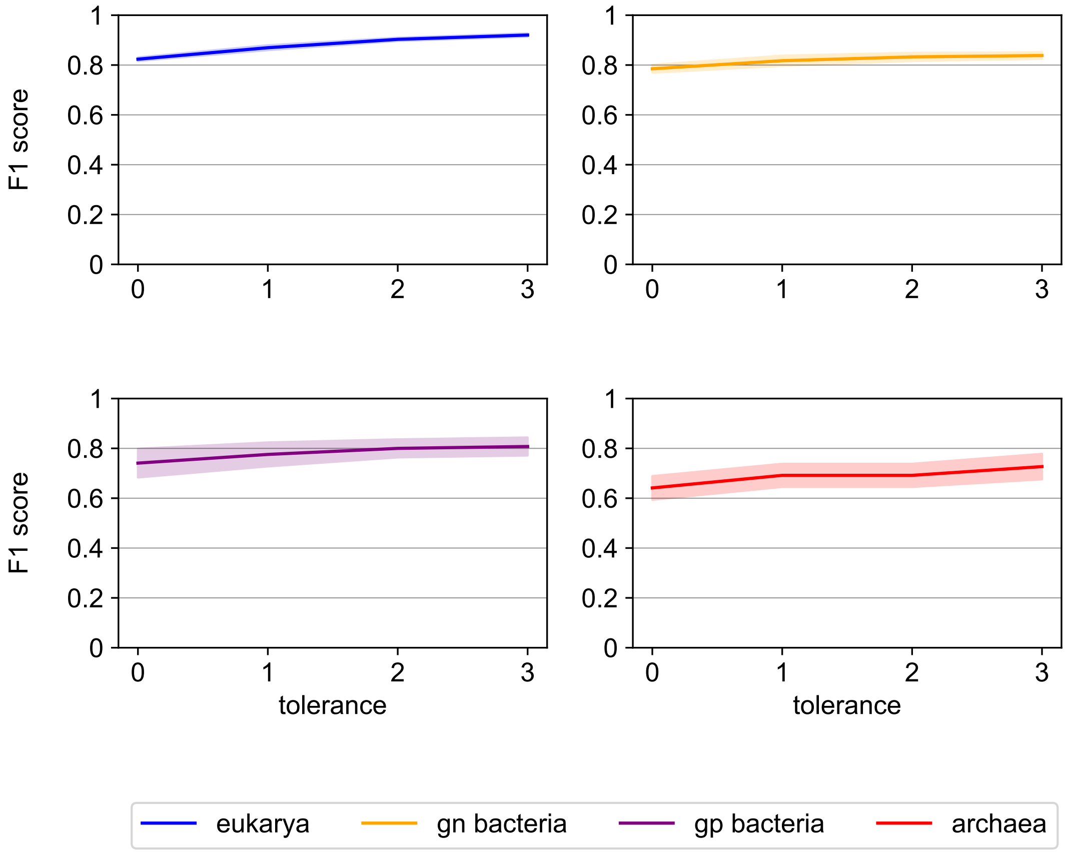


Figure 9: Performance of TSignal measured by the cleavage site F1 score over varying tolerance levels for Sec/SPase II and Sec/SPase IV. The F1 scores are computed over the whole dataset through nested cross-validation. The model is trained and tested five times, and the mean and 95% confidence estimates of the performance are plotted.

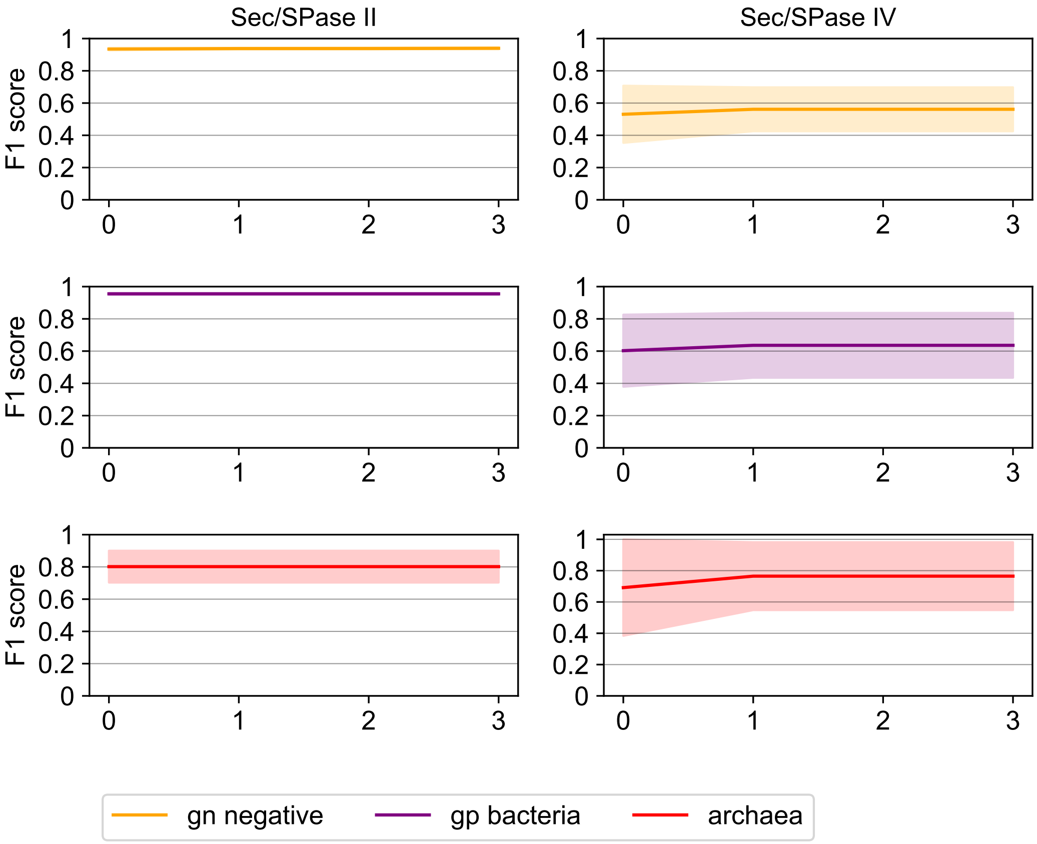


Figure 10: Performance of TSignal measured by the cleavage site F1 score over varying tolerance levels for Sec/SPase II and Sec/SPase IV. Noticeably, Sec/SPase II prediction performance is very similar across different tolerance levels, as the model easily learns to predict the CS on the Cys residue. Additionally, the Sec/SPase IV predictive performance is reasonably good and stable given the very low amount of data.

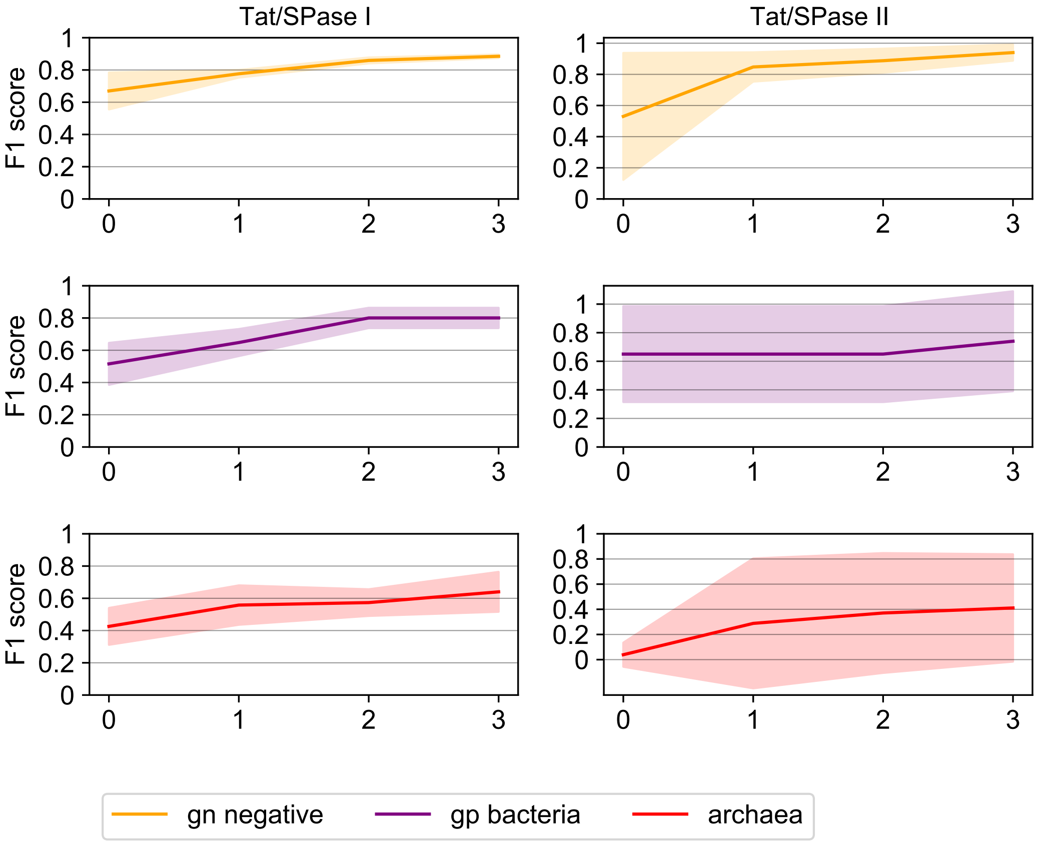


Figure 11: Performance of TSignal measured by the cleavage site F1 score over varying tolerance levels for Tat/SPase I and Tat/SPase II. The model is very good at distinguishing Tat from Sec directing SPs. Possibly due to the much longer SPs however, the cleavage sites in Tat/SPase I SPs seem to be less precise for tolerance 0 predictions, but become very good and stable for higher tolerance levels. In Tat/SPase II proteins, there is a noticeable increase in performance between tolerance 0 and 1 for gn-bacteria and archaea, due to the contextualized representation of residues. With only 19 sequences in the whole $\mathcal{D}$ dataset, gn-bacteria Tat/SPase II already have stable and good CS predictions, and we expect that archaea and gp-bacteria will also drastically increase with more data, as now only eight and six sequences respectively are present in all the available data $\mathcal{D}$.
